# Proteomic signaling of dual specificity phosphatase 4 (DUSP4) in Alzheimer’s disease

**DOI:** 10.1101/2023.09.13.557390

**Authors:** Erming Wang, Allen L. Pan, Pritha Bagchi, Srikant Ranjaraju, Nicholas T. Seyfried, Michelle E. Ehrlich, Stephen R. Salton, Bin Zhang

## Abstract

DUSP4 is a member of the DUSP (Dual-Specificity Phosphatase) subfamily that is selective to the mitogen-activated protein kinases (MAPK) and has been implicated in a range of biological processes and functions in Alzheimer’s disease (AD). In this study, we utilized stereotactic delivery of adeno-associated virus (AAV)-DUSP4 to overexpress DUSP4 in the dorsal hippocampus of 5×FAD and wildtype (WT) mice, then used mass spectrometry (MS)-based proteomics along with label-free quantification to profile the proteome and phosphoproteome in the hippocampus. We identified patterns of protein expression and phosphorylation that are modulated in 5×FAD mice and examined the sex-specific impact of DUSP4 overexpression on the 5×FAD proteome/phosphoproteome. In 5×FAD mice, a substantial number of proteins were up– or down-regulated in both male and female mice in comparison to age and sex-matched WT mice, many of which are involved in AD-related biological processes, such as the activated immune response or suppression of synaptic activities. Upon DUSP4 overexpression, significantly regulated proteins were found in pathways that were suppressed, such as the immune response, in male 5×FAD mice. In contrast, such a shift was absent in female mice. For the phosphoproteome, we detected an array of phosphorylation sites that are regulated in 5×FAD compared to WT, and are modulated by DUSP4 overexpression in each sex. Interestingly, the changes in 5×FAD– and DUSP4-associated phosphorylation occurred in opposite directions. Strikingly, both the 5×FAD– and DUSP4-associated phosphorylation changes were found for the most part in neurons, and play key roles in neuronal processes and synaptic function. Site-centric pathway analysis revealed that both the 5×FAD– and DUSP4-associated phosphorylation sites were enriched for a number of kinase sets in female, but only a limited number of sets of kinases in male mice. Taken together, our results suggest that male and female 5×FAD mice respond to DUSP4 overexpression via shared and sex-specific molecular mechanisms, which might underly similar reductions in amyloid pathology in both sexes, while learning deficits were reduced in only females with DUSP4 overexpression. Finally, we validated our findings with the sex-specific AD-associated proteomes in human cohorts and further developed DUSP4-centric proteomic network models and signaling maps for each sex.

## Introduction

Alzheimer’s disease (AD) is the most prevalent neurodegenerative disease and the most extensively studied cause of dementia^1^. Pathologically, AD is characterized by and manifests with neurofibrillary tangles that are formed from the improperly processed phosphorylated tau proteins in the intracellular space, and the accumulation of amyloid beta plaques in the intercellular space^2,3^. These abnormal aggregates are associated with oxidative stress and inflammation^4^, resulting in microglial activation and neurodegeneration in the brain^5^, and further causing the impairment or even loss of normal cognitive function and memory as age advances. Age, apolipoprotein E ɛ4 (APOE) and sex are the three greatest risk factors for AD^6^. In fact, sex is an important variable for AD patient stratification and personalized treatment^7^ and a recent study of a large number of transcriptomes show profound sex-specific changes and network remodeling in AD^8^. Mechanistically, sex-specific differential response to AD might be caused by the sex-specific differential transcriptional response to AD pathology^9^. Despite extensive studies investigating risk factors and neuropathogenesis of AD, the etiology and molecular and cellular mechanisms underlying AD are still largely unknown^2,3^, and the majority of the experimental drugs tested for AD have failed without showing significant efficacy^10^.

Proteomics analyses have been utilized to investigate mechanisms underlying neurodegenerative diseases because alterations in protein expression correlate better with phenotypes than changes in RNA expression^11^. Protein expression can be regulated at multiple levels, including transcriptional and epigenetic control over gene activity, and posttranscriptional modulation of RNA splicing, stability, and transport ^12^. Comparison of transcriptomic and proteomic profiling has revealed that ∼40% of the variation in protein expression is likely caused and regulated by posttranscriptional and translational/post-translational mechanisms^13,14^. Post-translational modifications (PTMs) regulate protein trafficking, function, and degradation, and thus aberrant PTMs of disease-relevant proteins would trigger abnormal alterations in pathological pathways, leading to disease progression^10^. Many studies of neurodegenerative diseases including AD have characterized PTMs of disease-relevant proteins such as tau^15^ and TDP-43^16^. Globally, MS-based proteomic analysis using both label-free^17^ and TMT-labeled^13,18^ approaches plus various enrichment strategies has emerged as an important paradigm to survey changes in the PTMs of AD patients and healthy controls. The results from these comprehensive surveys^13,17,18^ on PTMs provide valuable insight into the biochemical signaling pathways that drive AD pathogenesis and progression^15^.

Dual-specificity phosphatases (DUSPs) are a protein phosphatase subfamily with selectivity towards mitogen-activated protein (MAP) kinases^19^. DUSP4, a member of this family, has been shown to dephosphorylate MAPKs including ERK, JNK, and p38 kinases. In human epileptic brains, DUSP4 appears to function as a feedback inhibitor of pro-epileptogenic MAPK signaling^20^. Mechanistically, DUSP4 was demonstrated to be in the PRMT1-DUSP4-p38 axis to modulate cell differentiation^21^. DUSPs including DUSP4 have become an important focus of research in neurodegenerative diseases because of their identified contributions to many important biological processes, including neuroprotection, differentiation, and inflammation^19^. In our recent study^22^, we investigated the roles of DUSP4 and its downstream network in the development of learning behavior impairment and neuropathology in the 5×FAD amyloidopathy mouse model. We found that overexpression of DUSP4 improves learning behavior only in female 5×FAD, whereas β-amyloid load is reduced in both male and female mice^22^. Transcriptomics profiling plus pathway enrichment analysis further supported that DUSP4 may modulate AD phenotype in a sex-specific manner^22^.

In the present study, we sought to perform proteomics and phosphoproteomics analyses of the 5×FAD mice with and without DUSP4 overexpression, to identify proteins and phosphorylation modulated by DUSP4. We further compared our DUSP4-modulated proteomes with AD-associated protein signatures and networks derived from large proteomics studies of human postmortem brains with AD to understand how DUSP4 may contribute to AD pathogenesis. Finally, we developed sex-specific, DUSP4-centric proteomic network models and signaling maps.

## Results

We performed both proteomic and phosphoproteomic analyses using the label-free quantification of MaxQuant^23–25^ to analyze the mouse brain hippocampal samples extracted from 4 experimental groups that had been administered AAV-DUSP4 or AAV-GFP into dHc: 5×FAD-DUSP4 (n=7 females, n=4 males), 5×FAD-GFP (n=7 females, n=5 males), WT-GFP (n=5 females, n=7 males), and WT-DUSP4 (n=5 females, n=6 males) (Figure 1A, see Methods and Supplementary Data 1). As a quality control (QC), we verified the genotypes of mice by the western blot analysis using the antibody to transgenic human APP (6E10) and by microscopic observation of the GFP protein activity/fluorescence (see Methods). Based on this analysis, a male mouse originally identified as 5×FAD-GFP was re-classified as WT-GFP, and the downstream analysis was corrected. In addition, we conducted QC on the proteomic and phosphoproteomic data for further downstream processing (see Discussion).

**Figure 1.**
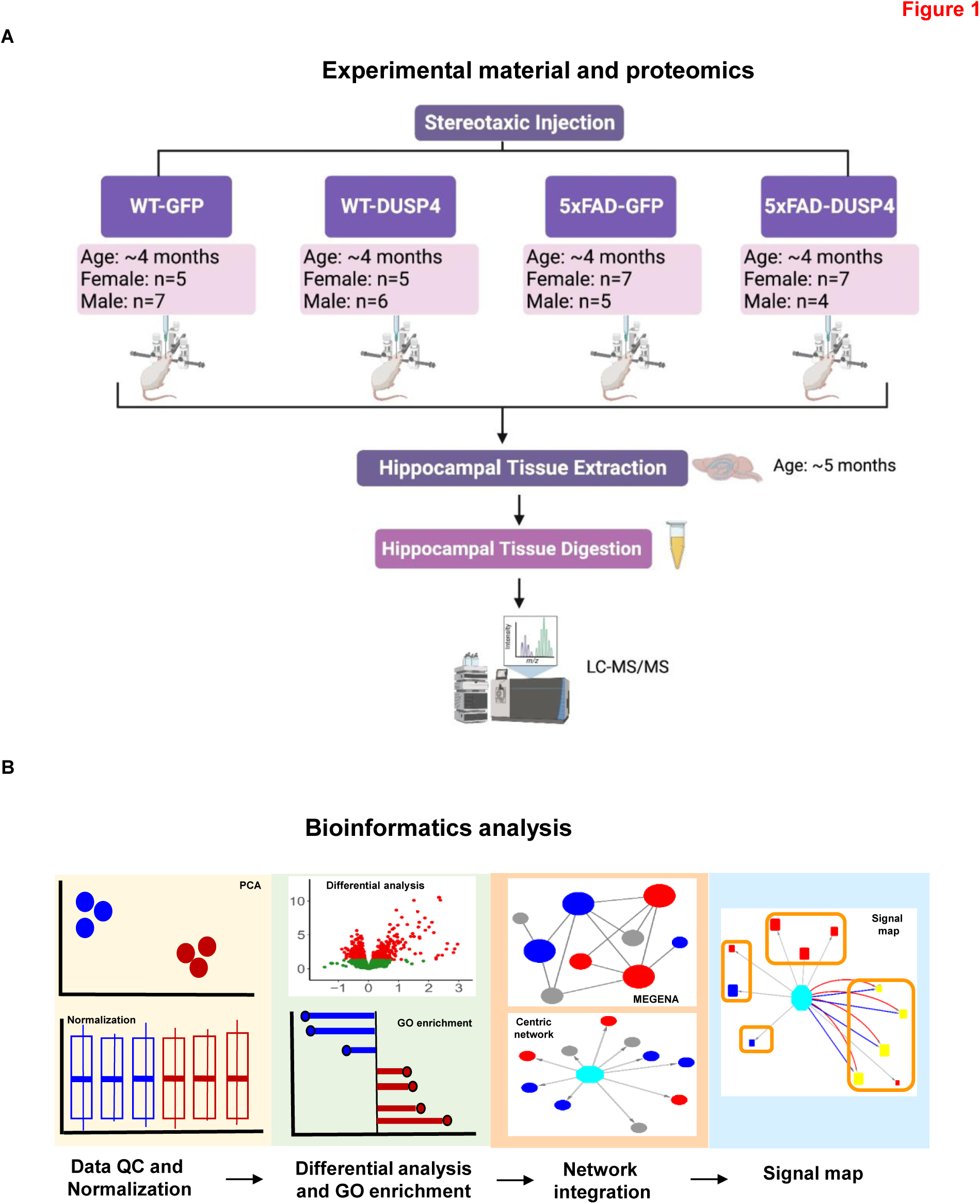
Schematic representation of experimental materials, data collection and bioinformatics workflow. (A) Experimental materials and data collection, created with BioRender.com. The female and male 5×FAD and wild type (WT) mice were injected with AAV5-DUSP4 or AAV5-GFP (control) into the hippocampus to over-express DUSP4 or GFP at 4 months of age. The hippocampal tissues were then extracted from 5×FAD and WT over-expressing DUSP4 or GFP one month after the surgery for subsequent LC-MS/MS analyses. (B) Downstream data processing workflow. The proteome and phosphoproteome data were first subject to quality control (QC) and normalization, followed by differential expression analysis to identify patterns of change under comparisons, which were further used to query gene ontology (GO) database for biological pathways and functional processes involved. Next, significant patterns of change in the mouse proteome and phosphoproteome were projected onto the human networks to find their relevance in human AD, and construction of sex-specific gene-centric networks and signal maps. MEGENA, Multiscale Embedded Gene Co-expression Network Analysis; PCA, principal component analysis.

In the present study, we focused our analysis on the two most critical comparisons, i.e., 5×FAD-GFP vs WT-GFP to identify proteins and phosphoproteins that are regulated in the 5×FAD mouse model compared to WT, and in comparisons of 5×FAD-DUSP4 to 5×FAD-GFP, to investigate the impact of DUSP4 overexpression on the proteome/phosphoproteome in 5×FAD. To simplify the presentation, we termed the comparison 5×FAD-GFP vs WT-GFP as 5×FADvsWT, and 5×FAD-DUSP4 vs 5×FAD-GFP as 5×FAD-DUSP4vs5×FAD.

Furthermore, we used the nominal p < 0.05 as a cut-off to include the proteins/phosphoproteins that are regulated by DUSP4 overexpression. Our experimental validation of selected proteins and integration with human proteomics showed that this cut-off is an effective criterion to determine the proteomic/phosphoproteomic signatures that are regulated by DUSP4 (see Discussion). Figure 1B highlights the bioinformatics workflow for data analysis and integration.

## Substantial numbers of differentially expressed proteins (DEPs) were regulated in 5×FAD and by DUSP4 overexpression

Together, we quantified 4,459 distinct proteins over the 46 samples. After QC (see Methods), we obtained 3578 unique proteins. We performed DEP analysis to reveal the mouse proteome which is impacted by the 5×FAD transgene and by DUSP4 overexpression. We identified 685 and 564 DEPs comparing 5×FADvsWT for female and male mice, respectively (Figure 2A; Supplementary Figure 1A; Supplementary Data 2A, B). We detected more DEPs that were down-regulated than up-regulated in 5×FADvsWT for mice of both sexes (Supplementary Figure 2). As expected, amyloid precursor protein (APP) expression was substantially elevated in 5×FAD mice of each sex (Figure 2A; Supplementary Figure 1A; Supplementary Data 2A, B). In the comparison of 5×FAD-DUSP4vs5×FAD, we found 295 and 335 DEPs for female and male mice, respectively (Figure 2B; Supplementary Figure 1B; Supplementary Data 2C, D). In contrast to the comparison of 5×FADvsWT, we detected more up-regulated DEPs than down-regulated ones in 5×FAD-DUSP4vs5×FAD in each sex (Supplementary Figure 2). As anticipated, DUSP4 protein levels were markedly increased in 5×FAD-DUSP4vs5×FAD for both female (fold-change (FC) = 6.7, p = 0.05) and male (FC = 22.8, p = 4.7E-6) mice, respectively (Supplementary Data 2C-D). Note that the APP protein expression was not altered in 5×FAD-DUSP4vs5×FAD.

**Figure 2.**
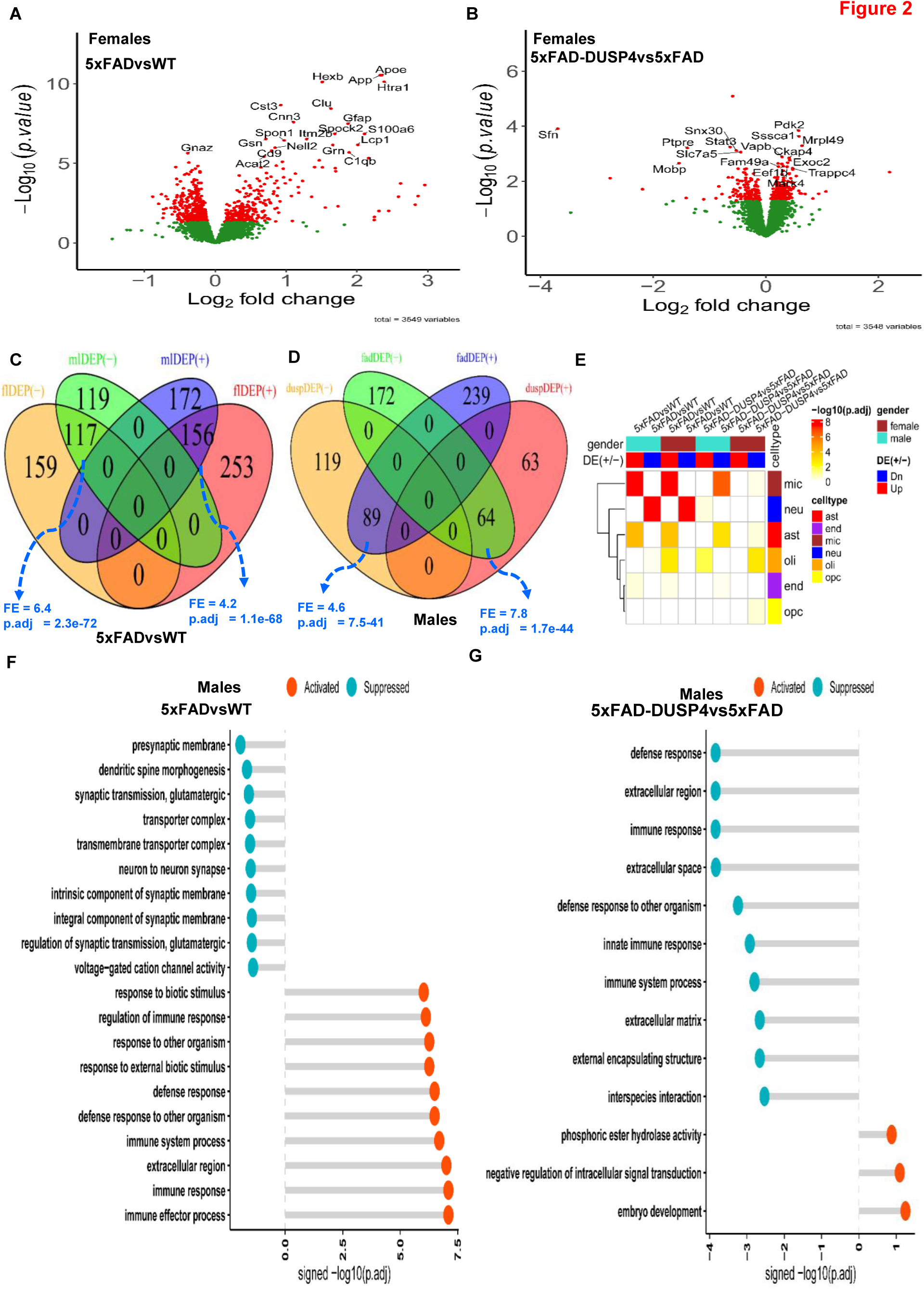
Analysis of differentially expressed proteins (DEPs). (**A**) Volcano plot showing the DEPs in 5×FADvsWT in female mice. (**B**) Volcano plot showing the DEPs in 5×FAD-DUSP4vs5×FAD female mice. In (**A**) and (**B**), each dot represents a protein, and highlighted are top-ranked DEPs. Dots in red are DEPs, whereas dots in green are not differentially expressed proteins. (**C**) Venn diagram showing the overlapping of DEPs in 5×FADvsWT between male and female mice. mlDEP(-) and mlDEP(+), are down– and up-regulated DEPs in male. flDEP(-) and flDEP(+), are down– and up-regulated DEPs in female. (**D**) Venn diagram showing the overlapping of DEPs between 5×FADvsWT and 5×FAD-DUSP4vs5×FAD in male mice. fadDEP(-) and fadDEP(+) are down– and up-regulated DEPs in 5XFAD. duspDEP(-) and duspDEP(+) are down– and up-regulated DEPs in DUSP4 overexpression. FE, fold enrichment. p.adj, BH-adjusted p value. (**E**) Enrichment of various mouse DEP lists in mouse cell-type signatures. The six mouse cell-type signatures were curated and described in^57^. (**G**) and (**F**) GO enrichment analysis on DEPs of 5×FADvsWT (**G**) and 5×FAD-DUSP4vs5×FAD (**F**) in male mice. x-axis, –log10(p.adj) split by enrichment groups, activated (positive) vs. suppressed (negative). y-axis, GO (gene ontology) terms.

We compared the DEP signatures across different comparisons for each sex. We separated up-regulated proteins from down-regulated ones to examine consistency in the directionality of protein expression changes. For each comparison, we observed significant overlap between the male and female DEP signatures in the direction of protein expression changes, and insignificant overlap in the opposite directions (Figure 2C; Supplementary Figure 3A). For example, the up–regulated signatures of males and females in 5×FADvsWT significantly overlap (fold enrichment (FE) = 4.2, FDR = 1.1E-68; Figure 2C) and the down-regulated signatures of males and females in 5×FAD-DUSP4vs5×FAD also significantly overlapped (FE = 1.8, FDR = 0.02; Supplementary Figure 3A). In contrast, in male mice, the up-regulated signature in 5×FAD-DUSP4vs5×FAD significantly overlap the down-regulated signature in 5×FADvsWT (FE = 7.8, FDR = 1.7E-44; Figure 2D). Similar results were observed in female mice (Supplementary Figure 3B and Supplementary Figure 2). These results show that DUSP4 overexpression reverse the abnormal proteomic changes in the 5×FAD mice in comparison with the wild type mice.

We further looked into the DEPs for cell-type specificity. We observed that the down-regulated signatures were enriched for the markers of neurons, whereas the up-regulated signatures were most enriched for the markers of microglia and astrocytes in 5×FADvsWT in both sex groups (Figure 2E), consistent with some previous finding of up-regulated immune response and neuronal damage, and down-regulated synaptic transmission ^26^. However, in 5×FAD-DUSP4vs5×FAD, we found that the down-regulated signatures were enriched for the markers of microglia and astrocytes in both sex groups, whereas the up-regulated signature in only males was enriched for the neuronal markers (Figure 2E). Thus, overexpression of DUSP4 affected all the major brain cell types, albeit with difference in enrichment significance across sex groups (Figure 2E).

We also examined biological pathways and functional processes in which these DEPs participated. In male 5×FAD mice, immune and defense response was activated while neuronal and synaptic functions were suppressed (Figure 2F). Similar results were observed for female 5×FAD mice (Supplementary Figure 4A). We then examined the effect of DUSP4 overexpression in 5×FAD mice. In male mice, DUSP4 overexpression activated pathways like intracellular signal transduction while suppressed immune and defense responses which were activated in 5×FAD mice (Figure 2G). However, in females DUSP4 overexpression affected a different set of pathways (Supplementary Figure 4B). Note that many pathways suppressed by DUSP4 overexpression in female mice (e.g., apoptotic process) are detrimental to cell functions (Supplementary Figure 4B). These results revealed sex-specific functions of DUSP4.

## DUSP4 overexpression caused significant changes in differentially expressed posttranslational modification (DEPTM) sites

We preprocessed the mass spectrometry (MS)-based phosphoproteome profiling using the R package PhosPiR, which removed MaxQuant-marked reverse sequences and potential contaminants, and have summarized the intensities for each phosphosite entry, termed PTM site (see Methods). The expression level (intensity) at each PTM site was obtained from quantile normalization and low-rank approximation imputation^27^. We removed any PTM site that had no gene name or PTM position information. The expression was further log2-transformed for the downstream analysis.

We obtained 7,124 distinct PTMs across the 46 samples, which spanned 2,222 unique proteins, averaging about 3 PTM sites per protein. We performed differential expression analysis on all the PTMs. We identified 982 and 557 DEPTMs in 5×FADvsWT for female and male mice, respectively (Figure 3A; Supplementary Figure 5A; Supplementary Data 3A, B). We detected more DEPTMs that were up-regulated than down-regulated in 5×FADvsWT in both sex groups (Supplementary Figure 6). In the comparison of 5×FAD-DUSP4vs5×FAD, we found 409 and 425 DEPTMs for female and male mice, respectively (Figure 3B; Supplementary Figure 5B; Supplementary data 3C, D). In contrast to the comparison of 5×FADvsWT, we detected more down-regulated than up-regulated DEPTMs in 5×FAD-DUSP4vs5×FAD for mice of either sex (Supplementary Figure 6). We then compared the DEPTM signatures across different comparisons in each sex in the same way as we conducted the DEP analysis (see above). Overall, a similar trend was observed for the DEPTMs as for the DEPs (Figure 3C, 3D; Supplementary Figure 7). In each comparison (5×FADvsWT or 5×FAD-DUSP4vs5×FAD), female and male mice shared a significant portion of DEPTMs with the same directionality, whereas in each sex, 5×FADvsWT and 5×FAD-DUSP4vs5×FAD showed significant overlap between their DEPTMs but with opposite directionality (Figure 3C, 3D; Supplementary Figure 7). These results again suggested DUSP4 overexpression might reverse the effects of the 5×FAD transgene on mice at the phosphoproteome level.

**Figure 3.**
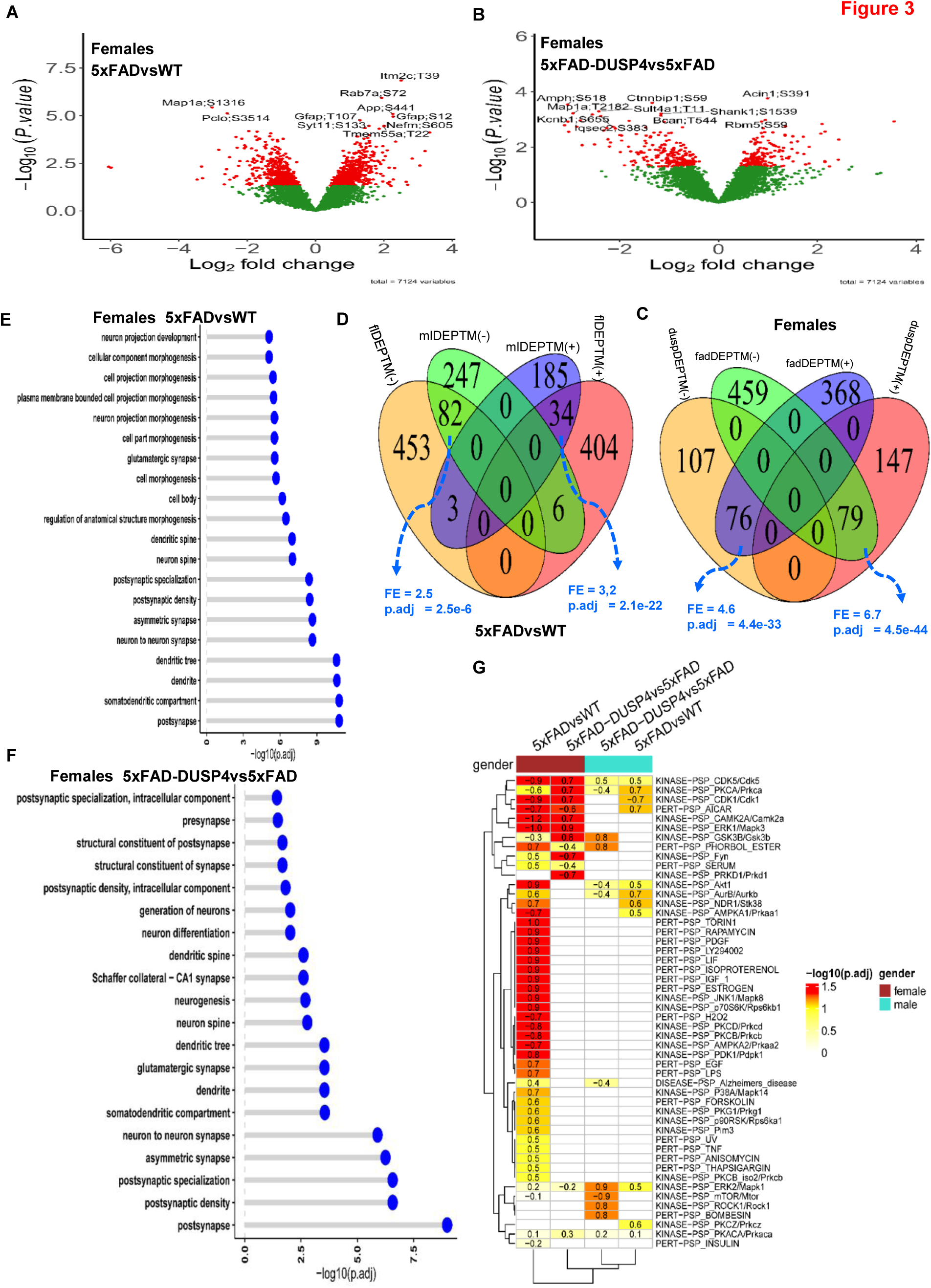
Analysis of differentially expressed PTM (DEPTM). (**A**) and (**B**) Volcano plots visualizing DEPTMs in 5×FADvsWT (**A**) and 5×FAD-DUSP4vs5×FAD (**B**) in female mice, respectively. In (**A)** and (**B)**, each dot represents a protein, and highlighted are top-ranked DEPTMs. Dots in red are DEPTMs, whereas dots in green are not differentially expressed PTMs. (**C**) Venn diagram showing the overlapping of DEPTMs between 5×FADvsWT and 5×FAD-DUSP4vs5×FAD in female mice. fadDEPTM(-) and fadDEPTM(+) are down– and up-regulated DEPTMs in 5XFAD. duspDEPTM(-) and duspDEPTM(+) are down– and up-regulated DEPTMs in DUSP4 overexpression. (**D**) Venn diagram showing the overlapping of DEPTMs in 5×FADvsWT between male and female mice. mlDEPTM(-) and mlDEPTM(+), are down– and up-regulated DEPTMs in male. flDEPTM(-) and flDEPTM(+), are down– and up-regulated DEPTMs in female. FE, fold enrichment. p.adj, BH-adjusted p value. (**E**) and (**F**) GO enrichment analysis on the DEPTMs of 5×FADvsWT (**E**) and 5×FAD-DUSP4vs5×FAD (**F**) in female mice. y-axis, GO terms. x-axis, –log10(p.adj). (**G**) PTM site enrichment analysis on various DEPTM signatures. Highlighted numbers are the score of fold enrichment.

We further explored the pathways in which the DEPTMs were involved. Since proteins may possess multiple sites of phosphorylation, we collapsed the DEPTM sites onto their respective protein levels. We define a differentially phosphorylated protein (DPP) as the one that contains at least one DEPTM. We obtained 665 and 418 DPPs in 5×FADvsWT for female and male mice, respectively, and 327 and 340 DPPs in 5×FAD-DUSP4vs5×FAD for female and male mice, respectively. As shown in Figure 3E, 3F, the most affected pathways are involved in neuronal processes and synaptic function for the DPPs (DEPTMs) across the comparisons in each sex (Supplementary Figure 8A, B), suggesting that both 5×FAD and DUSP4 might often influence the phosphorylation state of the proteins that are relevant to neuronal and synaptic function.

To more deeply delve into the signals represented in our phosphoproteome profiling, we applied the site-centric pathway analysis^28^ on our PTMs via the algorithm as described in the R package GSVA^29^ (see Methods). We examined how the PTMs are enriched for the PTM site-specific phosphorylation signatures^28^ (PTMsigDB). As shown in Figure 3G, in female 5×FADvsWT, the PTMs were enriched over more than half of the PTM sets in the mouse PTMsigDB. The top-ranked kinase PTM sets are KINASE-PSP_CAMK2A/Camk2a, KINASE-PSP_ERK1/Mapk3, KINASE-PSP_JNK1/Mapk8^30^ (Figure 3G), which are critical in AD neuropathogenesis. Strikingly, the PTMs from 5×FAD-DUSP4vs5×FAD in female mice are enriched for the PTM sets in the mouse PTMsigDB yet with an opposite directionality in enrichment score (ES) (Figure 3G), highlighting that DUSP4 overexpression altered the 5×FAD effects (activated or suppressed) on the PTM sets but in opposite directionality as 5×FAD in WT mice. In contrast, in male mice, the enrichment of PTMs in the mouse PTMsigDB was not very evident in spite of the enrichment in a few of PTM sets (Figure 3G). These results further suggested that DUSP4 overexpression might counteract the effects of 5×FAD transgene in mice in PTM site-centric pathways.

## DUSP4 overexpression resulted in reduction in STAT3 in 5×FAD mice

Hippocampal STAT3, human APP (hAPP), and DUSP4 protein levels were significantly altered in our proteomics data. To validate the changes of these proteins, western blotting was utilized to confirm protein levels. The results showed that hippocampal STAT3 protein levels were increased by about 110% in female 5×FAD mice overexpressing GFP (5×FAD-GFP), while male 5×FAD-GFP increased by about 65%, compared to age– and sex-matched wild type mice overexpressing GFP (WT-GFP) (Figure 4). STAT3 protein levels were reduced by about 65% in both female and male 5×FAD overexpressing DUSP4 (5×FAD-DUSP4) compared to age– and sex-matched 5×FAD-GFP (Figure 4). Although STAT3 protein levels were significantly reduced in female 5×FAD-DUSP4 compared to female 5×FAD-GFP, levels were significantly higher than female WT-GFP, while STAT3 protein levels in male 5×FAD-DUSP4 showed no significant differences compared to male WT-GFP (Figure 4). Western blot analyses confirmed the DUSP4 overexpression in both female and male mice administered with AAV-DUSP4. In addition, western blot analyses detected hAPP protein only in 5×FAD transgenic mice, which confirmed 5×FAD genotypes. These results validate the proteomics analyses.

**Figure 4.**
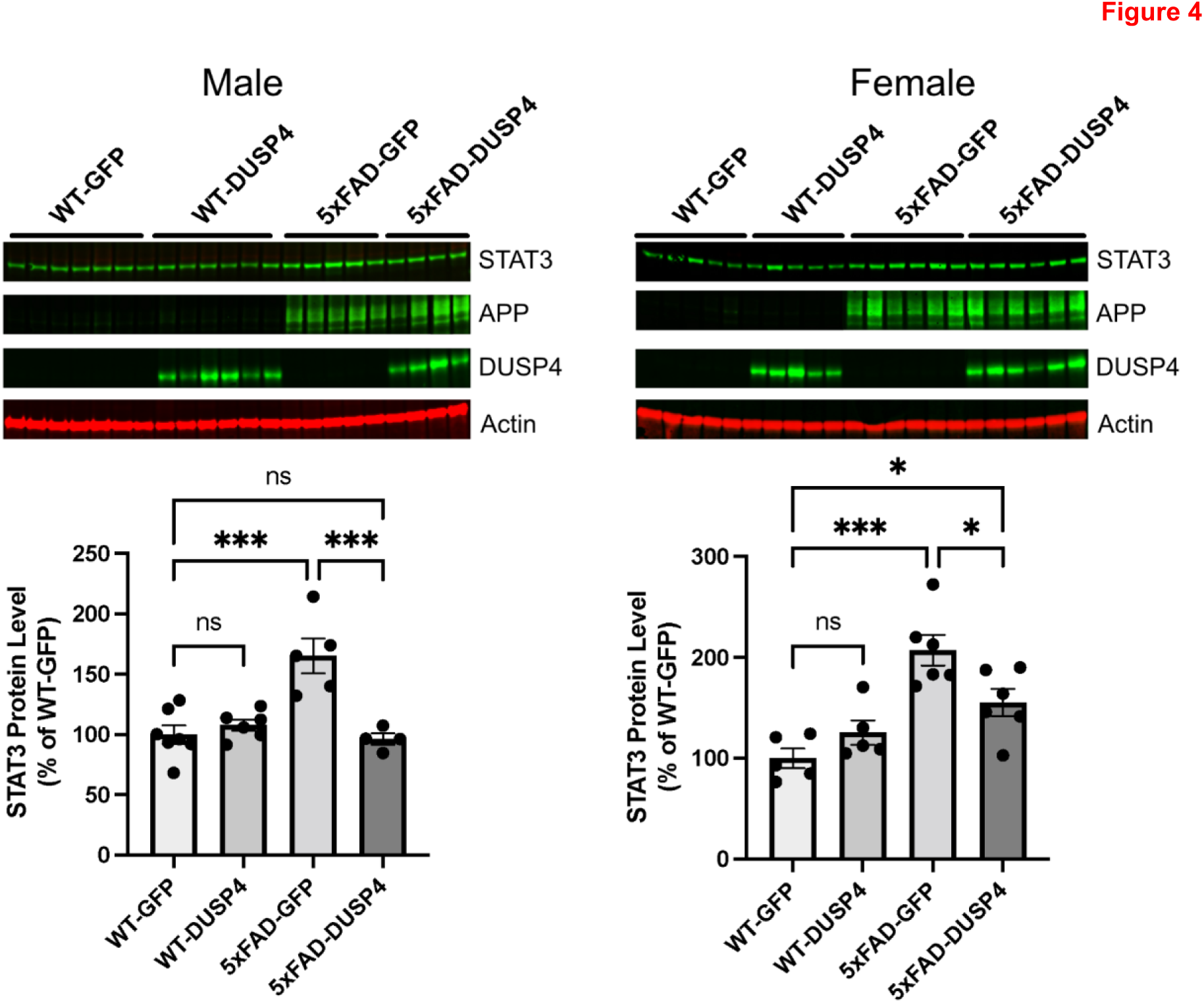
Western blot analyses of DEP from wild-type (WT) and 5×FAD mice overexpressing GFP or DUSP4. (A) Western blot analyses of hippocampal STAT3, human APP (hAPP), and DUSP4 protein levels in male WT and 5×FAD mice overexpressing GFP or DUSP4. n = 4-7 mice per group. (B) Western blot analyses of hippocampal STAT3, hAPP, and DUSP4 protein levels in female WT and 5×FAD mice overexpressing GFP or DUSP4. n = 5-6 mice per group. Error bars represent means ± SEM. Statistical analyses were performed using a One-way ANOVA followed by a Tukey’s post-hoc test, *p<0.05, ***p<0.001; ns = insignificant.

## The DUSP4 DEP and DEPTM signatures are enriched in human AD protein networks

We first compared the mouse DEP signatures in the present study with the human DEPs in AD that were derived from the proteomics profiling in the parahippocampal gyrus (PHG) of the MSBB cohort^31,32^. We stratified the human subjects over sex and thus obtained the sex-specific DEPs in AD vs normal healthy individual (NL) (Supplementary Data 4, and Methods). The mouse DEPs in 5×FADvsWT significantly overlapped the human DEP signatures, with the same directionality in both sexes though the overlap of male signatures was much less significant (Figure 5A, B). On the other hand, the DEP signatures in 5×FAD-DUSP4vs5×FAD in the male mice have marginally significant overlap with the human male DEP signatures with the opposite directions (Supplementary Figure 9A) while the signatures from the female mice don’t significantly overlap the respective human signatures (Supplementary Figure 9B). These results not only validated the mouse DEPs we identified but also suggested that our findings from the mouse proteomics might be relevant to human AD neuropathology.

**Figure 5.**
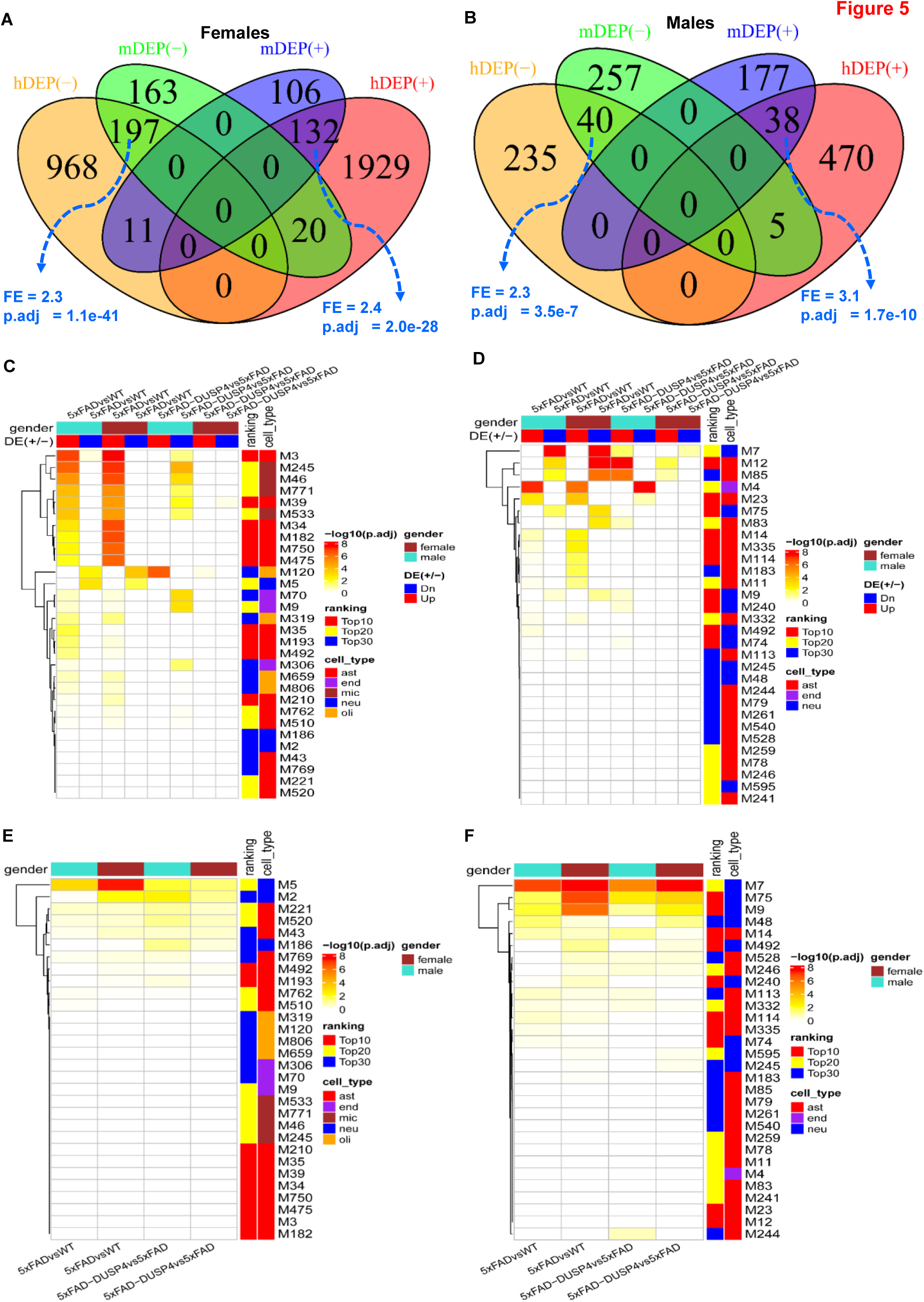
Integration of the DEPs and DEPTMs in mouse with the human co-expression networks. (**A**) Venn diagram revealing the overlap between female mouse DEPs in 5×FADvsWT and human female-specific DEPs in AD vs NL. (**B**) Venn diagram showing the overlap between male mouse DEPs in 5×FADvsWT and human male-specific DEPs in AD vs NL. In (A) and (B), mDEP(+) and mDEP(-), denote up– and down-regulated DEPs in mice, respectively, whereas hDEP(+) and hDEP(-), denote up– and down-regulated DEPs in human, respectively. (**C**) and (**D**) Heatmaps highlighting the enrichment of mouse DEP signatures over human proteomics MEGENA co-expression networks for the MSBB (C) and ROSMAP (D) cohorts. (**E**) and (**F**) Heatmaps highlighting the enrichment of mouse DEPTMs over human proteomics MEGENA co-expression networks for the MSBB (**E**) and ROSMAP (**F**) cohorts. DE(+/-), denotes up– or down-regulated DEPs, respectively. Ranking denotes the categories of the module ranking order in relevance to AD: Top10, Top20 and Top30 represent top-ranked 10, 20, and 30 AD modules, respectively. Cell-type, are the cell-type that is the most enriched for each module. ast, astrocytes; neu, neurons; endo, endothelial cells; mic, microglia; olig, oligodendrocytes. Because a protein may have more than one DEPTM site, we collapsed the DEPTMs to their respective protein levels, that is, a protein will represent all the DEPTMs that belong to it. Each field in the heatmap represents the intersection between a DEP or DEPTM signature over a module in the network. Only the 30 top-ranked AD-modules are shown.

We projected the mouse DEP signatures onto the MEGENA co-expression networks from the human proteomics^31^ to gain further understanding of their functional relevance to human AD. In the MSBB protein co-expression network, more than half (> 15) of the top 30 AD-associated modules were enriched for the mouse DEPs from 5×FADvsWT of both sexes (Figure 5C). The up-regulated DEPs in both male and female mice are enriched in the astrocyte (M3) and microglia modules (M245) while the down-regulated DEPs overlap significantly the neuronal modules (M5) (Figure 5C). We also observed the enrichment of the DEPs from 5×FAD-DUSP4vs5×FAD in the network, especially, the down-regulated DEPs in the male mice (Figure 5C). Similar results were found in the ROSMAP MEGENA network (Figure 5D). These results further validated the relevance of the mouse DEPs to human AD, and were consistent with the afore-described cell-type enrichment analysis (Figure 2E).

Furthermore, the mouse DEPTMs are also enriched in a number of top-ranked AD modules in the MSBB (Figure 5E) and ROSMAP (Figure 5F) protein co-expression networks. Importantly, the most enriched modules are neuron specific (M5 and M2 in the MSBB cohort, Figure 5E; M7 and M75 in the ROSMAP cohort, Figure 5F). These results are consistent with the previous pathway enrichment analysis (Figure 3E, 3F), and indicate that the DEPTMs are often involved in neuronal and synaptic functions.

## DUSP4 protein-centered networks are sex-specific

To formally identify the genes that are co-regulated with DUSP4 in AD, we leveraged a number of human AD cohorts as previously described ^33^ by examining the genes with significant correlations with DUSP4. We intersected the mouse DEP signatures in 5×FAD-DUSP4vs5×FAD with the human DUSP4-associated genes, and further constructed DUSP4 protein-centric networks for each sex (Figure 6A, 6B). There are more proteins positively correlated with DUSP4 protein/gene than those negatively correlated with DUSP4 protein/gene (Figure 6A, 6B). Impressively, the majority of the DUSP4-associated proteins are specifically expressed in either females (Figure 6A) or males (Figure 6B). Based on the sex-specific DUSP4-centric networks, we constructed the sex-specific DUSP4 signal maps (Figure 6C, 6D). As shown in Figure 6C, in females, DUSP4 is often involved in protein and lipid metabolism, in contrast to its involvement in synapse and myelin functions in males (Figure 6D). DUSP4 participates in endolysosomal pathways in both male and female mice, but in the opposite directions (Figure 6C, 6D). In summary, the results demonstrate that DUSP4 plays important roles in AD pathogenesis by regulating biological processes and functions shared by two sexes or distinct in each sex.

**Figure 6.**
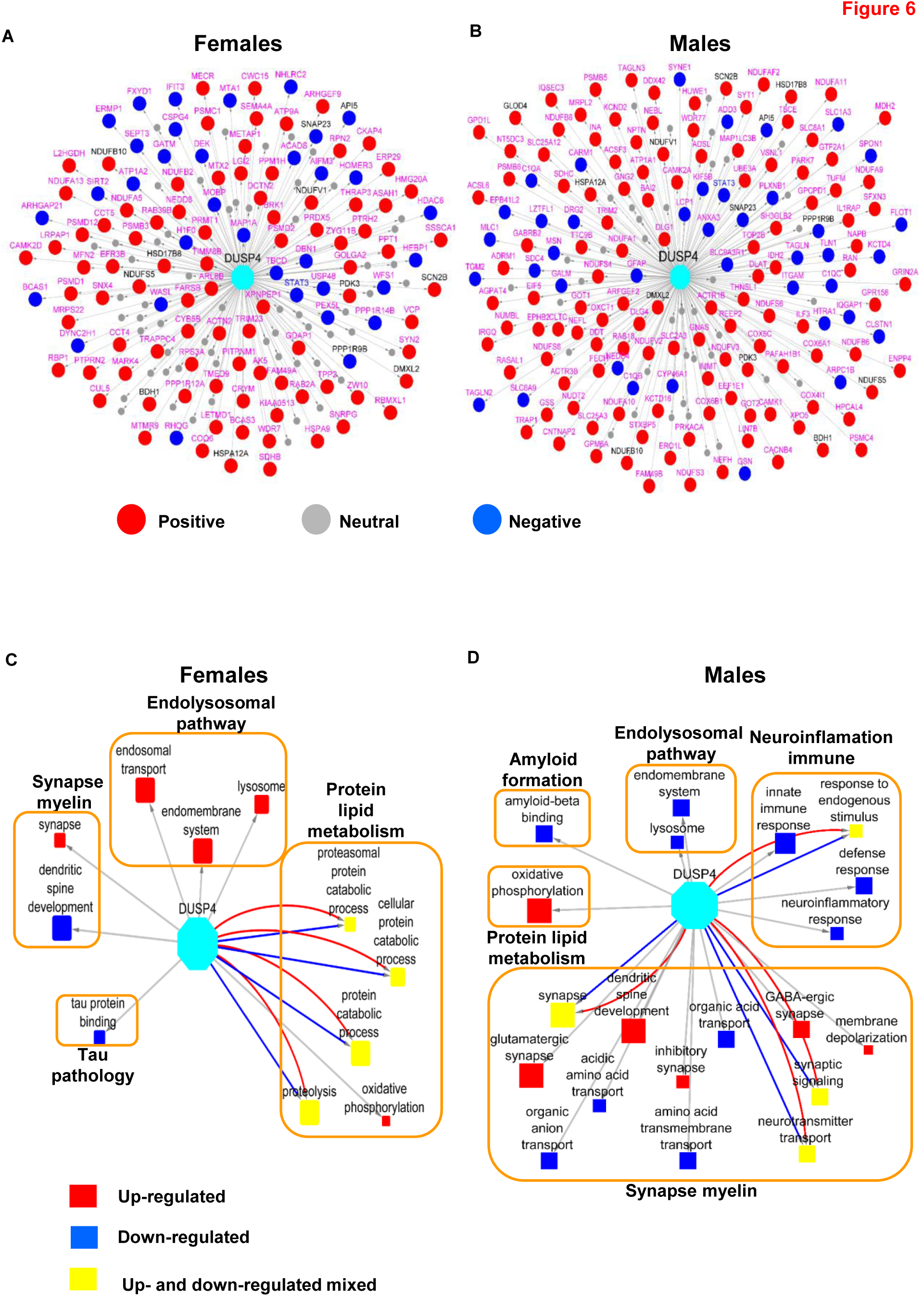
The DUSP4 protein-centric networks analysis. The networks were inferred using the DEP signatures from 5×FAD-DUSP4vs5×FAD for female (**A**) and male (**B**) mouse. Each node represents a gene (protein). Labeled in pink are the genes that are specifically associated with DUSP4 in female and male, respectively, whereas those in grey are common to both sexes. Red and blue nodes are positively– and negatively-associated with DUSP4, respectively, whereas grey nodes are not associated with DUSP4. (**C**) and (**D**) DUSP4 signal maps in female (**C**) and male (**D**), respectively, are shown. Each filled box denotes a GO term whose size is proportional to its enrichment for DUSP4-centric signatures. Red and blue highlight the enrichment only with positive and negative DUSP4-centric signatures, respectively, whereas yellow indicates the enrichment for both positive and negative DUSP4-centric signatures. The large unfilled boxes denote the parent categories of GO terms in AD.

## Discussion

In the present study, we investigated the proteins and phosphorylation sites that are modulated in 5×FAD mice, and examined the sex-specific impacts of DUSP4 overexpression on the 5×FAD proteome/phosphoproteome. In 5×FAD mice, a substantial number of proteins were up– or down-regulated in both male and female mice, and they were involved in AD-related biological processes, such as the activated immune response (upregulated in microglia and astrocytes) or suppressed synaptic activities (down-regulated in neurons). Upon DUSP4 overexpression, those dysregulated proteins and pathways (for example, immune response and defense response) were rescued. For the phosphoproteome, we detected an array of phosphorylation sites that are associated with 5×FAD and DUSP4 overexpression in each sex. However, the 5×FAD– and DUSP4-associated phosphorylation changes were in the opposite directions. Strikingly, both 5×FAD– and DUSP4-associated phosphorylation changes occurred mainly in neurons and these were predicted to regulate neuronal processes and synaptic function. Site-centric pathway analysis revealed that both the 5×FAD– and DUSP4-phosphorylation sites were enriched for a number of kinases in females, but only a limited number of kinases in male mice. Our study represents, to our knowledge, the first examination of the proteome and phosphoproteome that is modulated by DUSP4 and the determination of the significance of such modulation in AD.

DUSP4 is a regulator of the mitogen-activated protein kinase (MAPK) pathway, which regulates a wide variety of cellular signaling pathways, including stress responses, differentiation, and apoptosis^34^. Intriguingly, transcriptomic profiling of hippocampal RNAs in patients with Alzheimer’s disease (AD) showed a downregulation of DUSP4^35^, suggesting a potential role for DUSP4 in AD-associated pathogenesis. Our previous study in the 5×FAD AD animal model indicated that hippocampal DUSP4 overexpression rescued spatial memory deficits in female 5×FAD mice, but not in male 5×FAD mice^22^. In addition, transcriptomic profiling of 5×FAD mice overexpressing DUSP4 showed that differentially expressed genes (DEGs, false discovery rate (FDR) < 0.05) including Stat1, Stat2, and Ccl2 were downregulated in female 5×FAD-DUSP4 mice, while no DEGs (FDR<0.05) were detected in male 5×FAD overexpressing DUSP4. Furthermore, enrichment analysis of DEGs predicted that neuroinflammatory, interferon, and extracellular signal-regulated kinase (ERK)/MAPK signaling pathways were regulated in female 5×FAD overexpressing DUSP4^22^. While these transcriptomic data suggested a role for DUSP4 in AD-associated neuroinflammation, it is not clear how DUSP4 downregulated neuroinflammatory pathways. Consistent with our transcriptomic profiling of groups of mice with the same genotypes and AAV treatments, in the present study we observed up-regulated STAT1 protein in 5×FAD, which is reported in human AD^36^, whereas DUPS4 overexpression in 5×FAD female mice down-regulated STAT1 protein expression (Supplementary Data 2). Similar results were found in male mice although the changes were not robust by comparison in terms of p values. STAT2 and CCL2 were not profiled in the present study because of either low abundance at the protein expression level or large variation in expression among the samples, which caused their exclusion from further analysis. Overall, we observed significant overlaps for the 5×FAD– and DUSP4-associated signatures of both protein and phosphoprotein in female and male mice (Figure 2D and 3C; Supplementary Figure 3B and 7A), but in the opposite directions, indicating that DUSP4 overexpression may reduce AD-related deficits by reversing the dysregulated genes/proteins in 5×FAD in comparison with WT in a sex-specific manner. Furthermore, there exist significant differences in the sex-specific DUSP4-centric networks and signal maps (Figure 6). Taken together, our results that demonstrate sex-specific differences in the response of the 5×FAD proteome and phosphoproteomce to DUSP4 overexpression further support previous observations that DUSP4 overexpression reduces amyloidopathy in both sexes but learning deficits only in female 5×FAD mice^22^.

Microglia-associated neuroinflammation is characteristic of AD-associated pathology and was reported to be regulated by the ERK/MAPK signaling pathway^30^. Quantitative proteomics analyses showed that ERK1 and ERK2 were upregulated in postmortem AD human brains, and phosphorylated ERK was also increased in isolated microglia from 5×FAD mice^30^. In addition, proteomics analyses of the hippocampus in 5×FAD mice have revealed several pro-inflammatory markers including STAT3^26^, which can promote microglia-dependent neuroinflammation. For example, it was previously shown that deletion of microglial STAT3 in mice prevented microglia-dependent neuroinflammation^37^. The ERK/MAPK signaling pathway is a critical regulator of pro-inflammatory microglial activation, and microglial activation has been suggested as a contributor to the progression of AD^38^. In the present study, we found that DUSP4 overexpression in 5×FAD mice caused a reduction in STAT3 protein levels in both sexes (Figure 4 and supplementary Data 2). We subsequently queried the protein and protein interaction (PPI) network^39^ in AD, and obtained a STAT3-subnetwork (Supplementary Figure 10). Impressively, the STAT3-subnetwork is enriched for a number of GO terms critical in AD, such as amyloid formation, tau pathology, neuroinflammation and synapse and myelin functions (Supplementary Figure 10). Thus, it can be speculated that STAT3 is the connection point through which DUSP4 exerts its effects on AD, which might be one of the mechanisms underlying DUSP4 functionality that is shared in male and female mice.

In the present study, we used the nominal p < 0.05 as the cut-off in order to maximize inclusion of proteins/phosphoproteins that are regulated by DUSP4 overexpression. First, we carefully followed the standard experimental protocols and data processing pipelines (see Methods). As an example, we performed principal component analysis (PCA) on the protein and phosphoprotein expression data (Supplementary Figure 11), which was encouraging as it indicates that in general, mouse samples can be grouped together concordant to their genotypes. Then, we inspected and validated some of the proteins that were known to be regulated by 5×FAD. For example, we observed the up-regulation of the APP^22,26^, APOE^26^ and STAT3^26,40^ proteins in the 5×FAD mice of either sex, which is not only consistent with previously reported studies but was further confirmed by our experimental validation (Figure 4) and the integration analysis with the human proteomics profiling (Figure 5). For DEPTMs, we observed the phosphorylation site (APP;S441) in the APP protein was significantly up-regulated in both male and female 5×FAD mice (Figure 3A; Supplementary Data 3A-B). APP serine 441 has been inferred to be phosphorylated by a combination of experimental and computational evidence^41^ (The mouse App entry P12023, UniProtKB at https://www.uniprot.org/). Strikingly, DUSP4 overexpression resulted in a decreased level of phosphorylation at this site (APP;S441) in 5×FAD mice of either sex (Supplementary data 3C-D). S441 is found within the E2 dimerization domain of APP (aa374-565) (The mouse App entry P12023, UniProtKB at https://www.uniprot.org/). Whether S441 phosphorylation modulates antiparallel App dimer formation, heparin binding, and/or binding with other App interactors is to our knowledge unknown, although protein phosphorylation has been reported to modulate APP interactions^42^ and to occur in the APP ectodomain^43^. Similarly, we also observed 26 PTMs in the tau protein (Mapt gene), some of which displayed significant association with 5×FAD or DUSP4 (Supplementary information; Supplementary Figure 12). As an additional evidence, in female 5×FAD mice, we observed high consistency between the DEPs from the present study and the DEGs from our previous work^22^ (Supplementary Figure 13, and Supplementary results and discussion). These results and evidence together support this cut-off (nominal p < 0.05) as an effective criterion in determining the protein/phosphoprotein signatures regulated by DUSP4, albeit we cannot rule out any exceptions due to false discovery.

In summary, we characterized DUSP4-associated proteome and phosphoproteome, and unraveled the shared and sex-specific molecular mechanisms by which DUSP4 functions in 5×FAD mice.

## Methods Animal Studies

5×FAD transgenic mice were obtained from Jackson Labs (Bar Harbor, ME; JAX#34840) and were maintained on a mixed B6/SJL genetic background as described ^44^. Male and female 5×FAD and wild-type (WT) at 4 months of age were stereotactically infused with 1.0 μL of Adeno-Associated Virus (AAV)5-GFP or AAV5-DUSP4 (4×10^12^vg/ml) into dorsal hippocampus (dHc) (AP = –2.0 mm, ML = ± 1.5 mm, and DV = –2.0 mm relative to Bregma) at a rate of 0.2 μL per minute. AAV5-GFP (control) and AAV5-mouse DUSP4 (VectorBuilder Inc., Chicago, IL; AAV-5’ITR-CAG-mDUSP4-WPRE-BGHpA-3’ITR) (AAV5 serotype/AAV2 genotype) were prepared by the Vector Core at the University of North Carolina at Chapel Hill. All mice (Supplementary data 1) were housed under standard conditions (12 hour light-dark cycle with *ad libitum* access to food and water). All experimental procedures were conducted in accordance with the NIH guidelines for animal research and were approved by the Institutional Animal Care and Use Committee (IACUC) at the Icahn School of Medicine at Mount Sinai (ISMMS).

## Hippocampal tissue Collection

One month after the stereotactic infusion, mice were sacrificed and perfused with 20 mL ice-cold phosphate buffered saline (PBS). The whole hippocampal tissues were extracted from both hemispheres of the brains through gross dissection. Then the tissues were rinsed with PBS prior to storing at –80°C.

## Western blotting

STAT3, APP, and DUSP4 protein levels were analyzed by western blot as described^22^. Briefly, equal amounts of protein (20 µg) from each sample were resolved by electrophoresis in precast 4-12% Bis-Tris gels (Bio-Rad) and transferred to polyvinylidene difluoride (PVDF) membrane using the iBlot system (Invitrogen). Membranes were then incubated in Odyssey blocking buffer (92760001, LI-COR, Lincoln, NE) for 1 hour at room temperature before incubation with the following primary antibodies in a mixture of blocking buffer (92760001, LI-COR, Lincoln, NE) and 0.1% Tween-20 at 4°C overnight: anti-DUSP4 (1:1,000, ab216576, Abcam, Waltham, Boston); anti-Aβ (1:1,000, 803001, Biolegend, San Diego, CA); or anti-actin (1:1,000, MAB1501, Millipore Sigma). In the second day, membranes were washed with 0.1% Tween-20 in phosphate buffered saline (PBS) solution, and then incubated with a mixture of secondary antibodies: goat anti-rabbit 800CW (1:15,000, LI-COR, Lincoln, NE) and goat anti-mouse 680LT (1:20,000, LI-COR, Lincoln, NE) in Odyssey blocking buffer with 0.01% sodium dodecyl sulfate (SDS) and 0.1% Tween-20 at room temperature for 1 hour. Then the membranes were washed with 0.1% Tween-20 in PBS followed by PBS. The membranes were analyzed using an Odyssey infrared imaging system (LI-COR, Lincoln, NE). Protein bands were quantified using Odyssey Imager analysis software and were normalized using actin as an internal loading control.

## Proteomics/phosphoproteimics sample process, phosphopeptide enrichment, LC-MS/MS and MaxQuant analysis

Please see Supplementary information for details.

## Sex-specific differentially expressed protein (DEP) analysis in mouse

We used the label-free quantification (LFQ) intensity as the abundance for individual protein groups, termed proteins throughout the present study. We next removed proteins that either are potential contaminants or contain no protein (reverse) or were only identified by site. We only retained proteins that have expression in more than half of the samples. We then log2-transformed the protein expression, and imputed the missing values via the function in the R package ‘impute’^45^ with the default parameters. Finally, we normalized the protein expression via median centering^46^. To identify DEPs, we performed a pair-wise comparison to detect significant changes in protein level between any two experimental mouse groups (Supplementary data 2) in each mouse sex by the moderated t-test implemented in the limma package^47,48^. DEPs were determined to have a nominal p value < 0.05 (see Discussion).

## Sex-specific differentially expressed posttranslational modification (DEPTM) analysis in mouse

We preprocessed the mass spectrometry (MS)-based phosphoproteome profiling using the R package PhosPiR, which remove MaxQuant-marked reverse sequences and potential contaminants and summarize the intensities for each phosphosite entry, termed PTM site (see Methods). The expression level (intensity) at each PTM site was obtained following quantile normalization and low-rank approximation imputation^27^. We removed any PTM site that has no gene name or PTM position information. The expression was further log2-transformed for the downstream analysis. To identify DEPTMs, we performed a pair-wise comparison to detect significant changes in PTM level between any two experimental mouse groups (Supplementary data 3) across mouse sex by the moderated t-test implemented in the limma package^47,48^. DEPTMs were determined to have a nominal p value < 0.05 (Discussion).

## Gene set variation analysis (GSVA) on PTM site enrichment analysis

We applied the site-centric pathway analysis^28^ on our PTMs via the algorithm as described in the R package GSVA^29^. We examined the PTMs for enrichment over the mouse database of PTM site-specific phosphorylation signatures (PTMsigDB)^28^. We used the PTM expression matrix as the input to calculate the enrichment score for the sets of PTM sites in the mouse PTMsigDB in each sample across sex by GSVA^29^. We then performed differential analysis on the enrichment scores using the limma package^47,48^, which was followed by multiple test adjustment by the Benjamini-Hochberg (BH) method.

## Gene ontology (GO) enrichment analysis, and plot visualization

We used the R package clusterProfiler ^49^ to identify biological processes that are up– and down-regulated in each comparison. For DEP enrichment analysis, we used the function gseGO^49^ since we want to capture both activated and suppressed enrichment, whereas we used enrichGO^49^ function for the enrichment of the protein signatures derived from DEPTMs. The plots were generated by using Cytoscape (3.7.2) and the R packages ComplexHeatmap^50^, ggplot2, ggpubr, EnhancedVolcano, and SuperExactTest^51^. The R version was 4.2.0.

## Sex-specific DEP analysis in human cohorts

We performed DEP analysis in AD vs NL over the proteomics profile in two human AD cohorts using the postmortem tissue from two different brain region: the parahippocampal gyrus (PHG) for the Mount Sinai Brain Bank (MSBB)^32^ and the prefrontal cortex (PFC) for the Religious Orders Study and Memory and Rush Aging (ROSMAP)^52,53^ cohort, respectively. The processing, normalization and co-variable adjustment for the human proteomics are as previously described ^31^. In the present study, we stratified the subjects by sex and then identified the sex-specific DEPs in AD in comparison with NL (Supplementary Data 4).

## Co-expression network analysis

Gene co-expression networks on the proteomes in the human cohorts were identified by using Multiscale Embedded GEne co-expression Network Analysis (MEGENA) as described in^31,54^.

## Construction of DUSP4 centric gene co-expression networks

We constructed DUSP4 centric consensus gene co-expression networks from 8 datasets from 3 cohorts including the MSBB (4 brain regions), ROSMAP (1 brain region) and HBTRC (3 brain regions) ^33^. In each dataset, the genes significantly correlated with DUSP4 (FDR < 0.05) were identified. From the significant correlations, a directional voting method was applied to calculate the frequency of negative or positive correlations between DUSP4 and each other gene. The DUSP4 centric network were thus defined as a function of frequency threshold n (=1, 2, …, 8}^33^. We then projected the DUSP4-associated DEPs in each sex onto the DUSP4 centric network, thus obtaining male or female-specific DUSP4-centric networks, respectively.

## Development of DUSP4 centric signaling maps

In our recent publication^55,56^, we identified about 70 AD-related GO terms including pathways related to Aβ, oxidative stress, tau NFT, and synaptic function. We used these gene sets to assess the relevance of a gene signature of interest with AD. Specifically, the connection between the gene signature of a target and an AD gene set is quantified by the enrichment score (–log10(FDR)), where FDR was determined by Fisher’s Exact Test and multiple testing correction. We only keep the connections with an FDR < 0.05, thus, the higher the score the more relevant a target to a pathway (i.e., GO term). All the significant connections constitute the target’s signaling map in AD.

## Availability of data and software code

The mouse proteomics and phosphoproteomics profiling data are available via syn52138250 at the AD Knowledge Portal (https://adknowledgeportal.synapse.org). The AD Knowledge Portal is a platform for accessing data, analyses, and tools generated by the Accelerating Medicines Partnership Alzheimer’s Disease (AMP-AD) Target Discovery Program and other NIA-supported programs to enable open-science practices and accelerate translational learning. The data, analyses, and tools are shared early in the research cycle without a publication embargo on secondary use. Data are available for general research use according to the following requirements for data access and data attribution (https://adknowledgeportal.synapse.org/DataAccess/Instructions). All codes are available up request.

## Competing interests

The authors declare that they have no competing interests. N.T.S. is co-founder of Emtherapro and Arc Proteomics.

## Funding

This work was supported in parts by grants from the National Institutes of Health (NIH)/National Institute on Aging (RF1AG054014 to B.Z., RO1AG068030 to B.Z., RF1AG074010 to B.Z., UO1AG046170 to B.Z., RF1AG057440 to B.Z., RO1AG057907 to B.Z., R01AG062355 to S.R.S., M.E.E., B.Z.; RF1AG062661 to S.R.S., M.E.E.; RF1-AG071587 to S.R., N.T.S.), the Cure Alzheimer’s Fund (to S.R.S., M.E.E.), and R01-NS114130 (S.R.).

## Authors’ contributions

Conceptualization, B.Z., S.R.S., M.E.E.; Investigation, E.W., A.L.P., P.B., N.T.S., S.R.S. and B.Z.; Resources, B.Z., S.R.S., M.E.E.; Writing – Original Draft, E.W., A.L.P., P.B.; Writing – Review & Editing, All Authors; Supervision, B.Z., S.R.S., M.E.E.; Funding Acquisition, S.R.S., B.Z., M.E.E..

## Acknowledgments

Dr. Minghui Wang is thanked for helping to download the proteomics and phosphoproteomics profiling data, and useful discussion during the manuscript preparation.

