## Supplementary information for "Proteomic signaling of dual specificity phosphatase 4 (DUSP4) in Alzheimer’s disease"

---

---

### Supplementary methods

#### Proteomics/phosphoproteimics sample preparation

Each tissue sample was added to 300 µL of urea lysis buffer (8 M urea, 10 mM Tris, 100 mM $NaH_2PO_4$, pH 8.5, including 3 µL (100x stock) HALT(-EDTA) protease and phosphatase inhibitor cocktail (Pierce)) in a 1.5 mL Rino tube (Next Advance) harboring stainless steel beads (0.9-2 mm in diameter). Samples were homogenized twice for 5-minute intervals in the cold room (4 °C). Protein homogenates were transferred to 1.5 mL Eppendorf tubes on ice and were sonicated (Sonic Dismembrator, Fisher Scientific) 3 times for 5 sec each with 5 sec intervals of rest at 30% amplitude to disrupt nucleic acids and were subsequently centrifuged at 4° C. Protein concentration was determined by the bicinchoninic acid (BCA) method, and samples were frozen in aliquots at -80 °C. Protein homogenates (500 µg) were treated with 5 mM dithiothreitol (DTT) at room temperature for 30 min, followed by 10 mM iodoacetimide at room temperature for 30 min in the dark. Protein samples were digested with 1:25 (w/w) lysyl endopeptidase (Wako) at room temperature overnight. Next day, samples were diluted with 50 mM $NH_4HCO_3$ to a final concentration of less than 2 M urea and were further digested overnight with 1:25 (w/w) trypsin (Promega) at room temperature. The resulting peptides were desalted with HLB column (Waters) and were dried under vacuum.

#### Phosphopeptide enrichment

Ten µg peptide was reserved for total proteome measurement and rest of the peptide was subjected to phosphopeptide enrichment using Thermo Scientific High-Select™ Fe-NTA Phosphopeptide Enrichment Kit. Dries peptides were submitted for mass spectrometry.

**LC-MS/MS analysis**

The data acquisition by LC-MS/MS was adapted from a published procedure [1]. Derived peptides were resuspended in the loading buffer (0.1% trifluoroacetic acid, TFA) and were separated on a Water's Charged Surface Hybrid (CSH) column (150 µm internal diameter (ID) x 15 cm; particle size: 1.7 µm). The samples were run on an EVOSEP liquid chromatography system using the 15 samples per day preset gradient (88 min) and were monitored on a Q-Exactive Plus Hybrid Quadrupole-Orbitrap Mass Spectrometer (ThermoFisher Scientific). The mass spectrometer cycle was programmed to collect one full MS scan followed by 20 data dependent MS/MS scans. The MS scans (400-1600 m/z range, $3 \times 10^6$ AGC target, 100 ms maximum ion time) were collected at a resolution of 70,000 at m/z 200 in profile mode. The HCD MS/MS spectra (1.6 m/z isolation width, 28% collision energy, $1 \times 10^5$ AGC target, 100 ms maximum ion time) were acquired at a resolution of 17,500 at m/z 200. Dynamic exclusion was set to exclude previously sequenced precursor ions for 30 seconds. Precursor ions with +1, and +7, +8 or higher charge states were excluded from sequencing.

**MaxQuant analysis**

Label-free quantification analysis was adapted from a published procedure[1]. Spectra were searched using the search engine Andromeda, integrated into MaxQuant, against 2020 Uniprot/Swiss-Prot mouse database (17,041target sequences). Serine, threonine, and tyrosine phosphorylation (+79.9663 Da), methionine oxidation (+15.9949 Da), asparagine and glutamine deamidation (+0.9840 Da), and protein N-terminal acetylation (+42.0106 Da) were variable modifications (up to 5 allowed per peptide); cysteine was assigned as a fixed carbamidomethyl modification (+57.0215 Da). Only fully tryptic peptides were considered with up to 2 missed cleavages in the database search. A precursor mass tolerance of ±20 ppm was applied prior to mass accuracy calibration and ±4.5 ppm after internal MaxQuant calibration. Other search settings included a maximum peptide mass of 6,000 Da, a minimum peptide length of 6 residues, 0.05 Da tolerance for orbitrap and 0.6 Da tolerance for ion trap MS/MS scans. The false discovery rate (FDR) for peptide spectral matches, proteins, and site decoy fraction were all set to 1 percent. Quantification settings were as follows: re-quantify with a second peak finding attempt after protein identification has completed; match MS1 peaks between runs; a 0.7 min retention time match window was used after an alignment function was found with a 20-minute RT search space. Quantitation of proteins was performed using summed peptide intensities given by MaxQuant. The quantitation method only considered razor plus unique peptides for protein level quantitation.

**DESeq2 analysis**

The RNA-seq raw read quality control and preprocessing are as described in [2]. Differentially expressed genes were identified using the R package DESeq2[3]. The input count matrix was the read counts summarized at the gene level per sample. Pre-filtering was used to screen out low count genes so that only genes with at least 2 reads, in total, across the samples in each comparison were kept. Differential expression analysis was performed using the DESeq function, followed by the log fold change shrinkage[4] via the type of apeglm. To be consistent with DEP identification, we defined differentially expressed genes (DEG) as those with a nominal $p < 0.05$. We further termed a DEG as up-regulated if its log2FoldChange > 0 in the comparison or down-regulated if its log2FoldChange < 0.

**Supplementary results and discussion**

**DUSP4- and 5xFAD-associated PTMs in the tau protein**

Another hallmark in AD neuropathology are the neurofibrillary tangles, which are caused by the abnormally phosphorylated tau protein[5,6]. Remarkably, we detected 26 PTM sites in the tau protein (Matp gene) across the samples (Supplementary Figure 12). Importantly, the majority of these PTMs are present in the database (UniProtKB at https://www.uniprot.org/). In addition, as shown in Supplementary Figure 12, some of the PTMs are significantly associated with DUSP4 or 5xFAD. Strikingly, again, most of the PTMs displayed an opposite directionality in their association with DUSP4 vs. 5xFAD in each sex (Supplementary Figure 12). For example, in male mice, the PTM (Mapt;S554) is up-regulated by DUSP OE, whereas it is down-regulated by 5xFAD (Supplementary Figure 12). Once again, these results suggested that DUSP4 OE might reverse the effects of the 5xFAD transgene on the phosphorylation of tau protein in mouse brain.

**DEG and DEP signatures are often overlapped**

In our previous work, we reported DUSP4- or 5xFAD-associated genes (mRNAs) [2]. To gain insight into the consistency between the DEGs and DEPs, we inspected the overlap between the DEG and DEP signatures over each comparison in each sex. For 5xFADvsWT in female mice, DEGs and DEPs are overlapped significantly and in the same directionality (Supplementary Figure 13). For example, the up-regulated DEG (mDEG(+)) is highly significantly overlapped with the up-regulated DEP (mDEP(+)) signature (FE = 3.0, p.adj = 9.0e-47, Supplementary Figure 13), highlighting the concordance in change in expression between the mRNAs and proteins caused by 5xFAD.

In 5xFAD-DUSP4vs5xFAD, down-regulated DEG and DEP signatures were significantly overlapped in female (Supplementary Figure 14A) and male mice (Supplementary Figure 14B). However, an insignificant overlap was found for the up-regulated DEG and DEP signatures albeit we observed > 10 hits in overlap for each sex (p.adj is close to 0.2, Supplementary Figure 14). In contrast, for 5xFADvsWT in male mice, we did not observe significant overlaps between DEG and DEP signatures (data not shown). This might be due to that the RNA-seq assay is not sensitive enough to detect DEGs for this comparison in male at the time when the sampling took place or else any biological mechanism(s) underlying this discrepancy. Clearly, more future research is needed to clarify such inconsistency.

**Supplementary Figures legends**

**A**

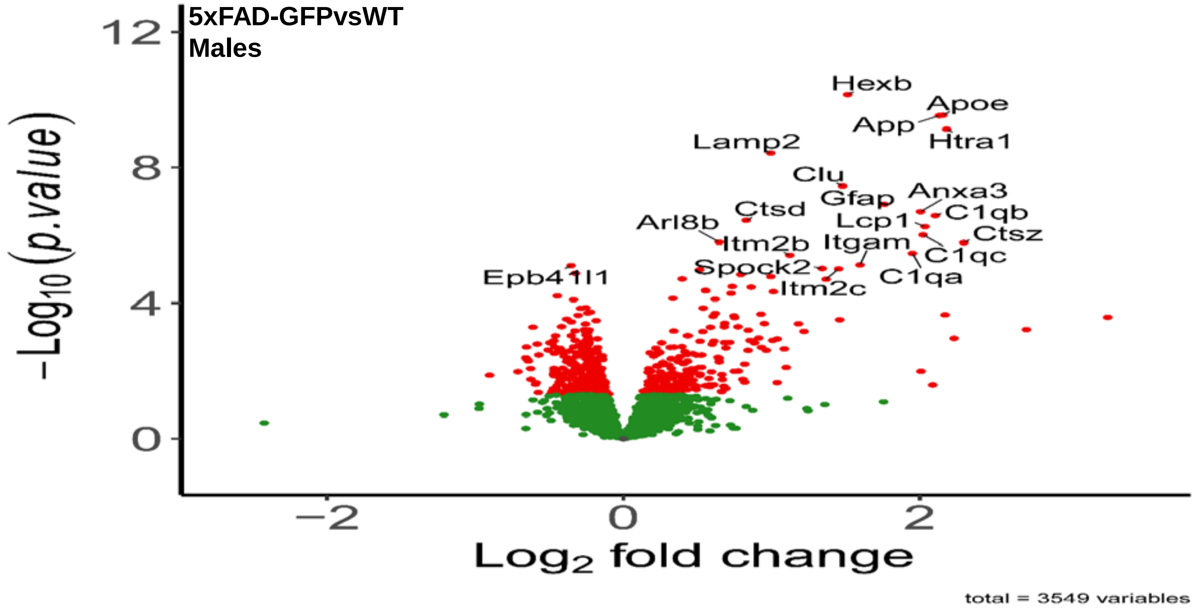
5xFAD-GFPvsWT
Males
Hexb
Apoe
App
Htra1
Lamp2
Clu
Gfap
Anxa3
Ctsd
Lcp1
C1qb
Arl8b
Ctsz
Itm2b
Itgam
C1qc
Spock2
C1qa
Epb41l1
Itm2c
12
8
4
0
−2
0
2
$-\text{Log}_{10}(p.value)$
$\text{Log}_2$ fold change
total = 3549 variables

**B**

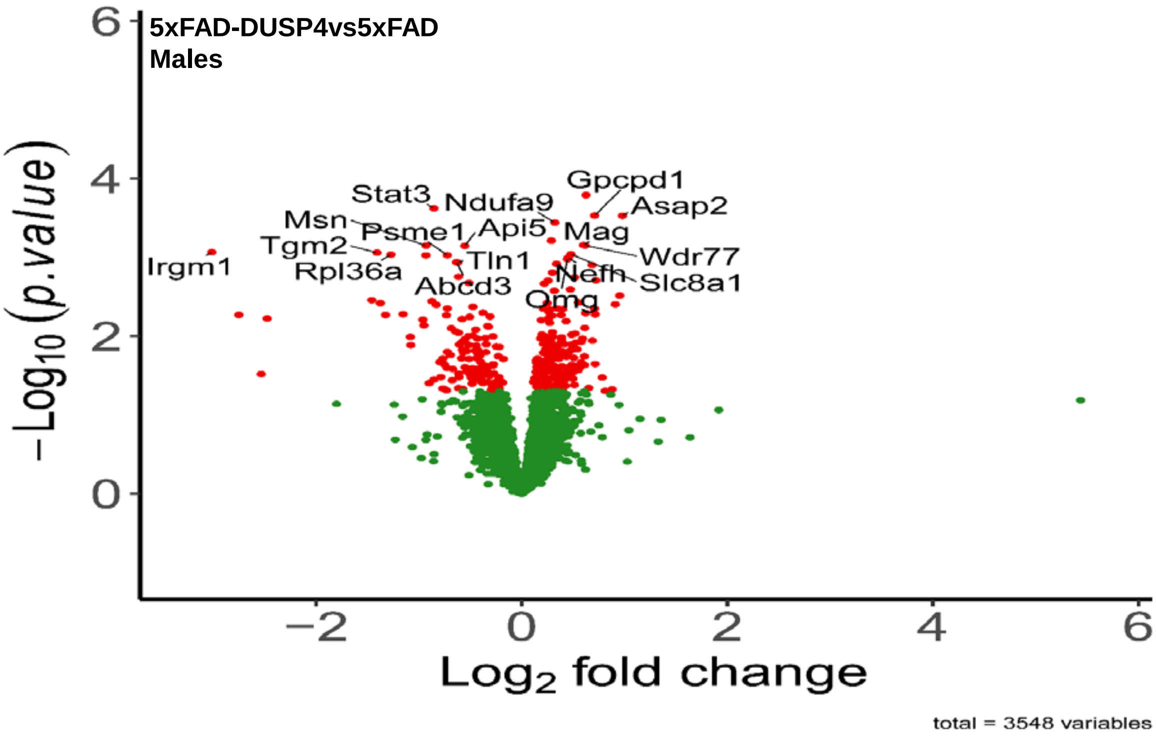
5xFAD-DUSP4vs5xFAD
Males
Gpcpd1
Stat3
Ndufa9
Asap2
Msn
Psme1
Api5
Mag
Tgm2
Wdr77
Irgm1
Tln1
Rpl36a
Nefh
Slc8a1
Abcd3
Omg
6
4
2
0
−2
0
2
4
6
$-\text{Log}_{10}(p.value)$
$\text{Log}_2$ fold change
total = 3548 variables

**Supplementary Figure 1. Volcano plots for DEP analysis.** (**A**) The DEPs in 5xFADvsWT in the male mice. (**B**) The DEPs in 5xFAD-DUSP4vs5xFAD in the male mice. Each dot represents a protein. Highlighted are top-ranked DEPs. Dots in red are DEPs, whereas dots in green are not differentially expressed proteins.

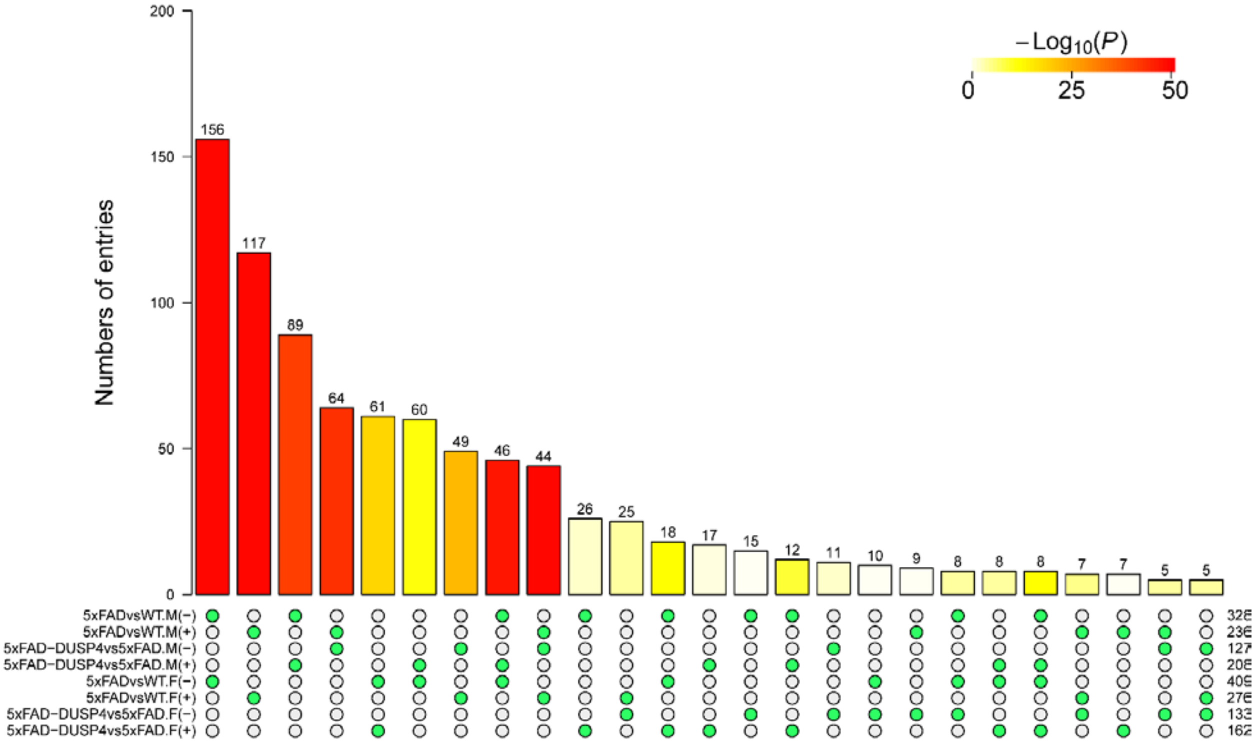

**Supplementary Figure 2. Summary of the overlap and significance of the DEP signatures via the superExactTest.** On the bottom of the bar graph, each row represents a DEP signature split by directionality in change in expression: - and +, down- and up-regulated, respectively. F and M, female and male, respectively. For example, 5xFADvsWT.M(-) stands for the down-regulated DEP signature in 5xFADvsWT in male mice. On the right are the numbers of the DEPs in each signature. Only shown here are the overlaps up to 3 groups. The number at the top of each bar is the number of DEPs overlapped among the groups (filled green circles) across rows under each bar. The color in each bar is proportional to the significance of the overlap (-log10(P)).

**A** **Male vs female in 5xFAD-DUSP4vs5xFAD**

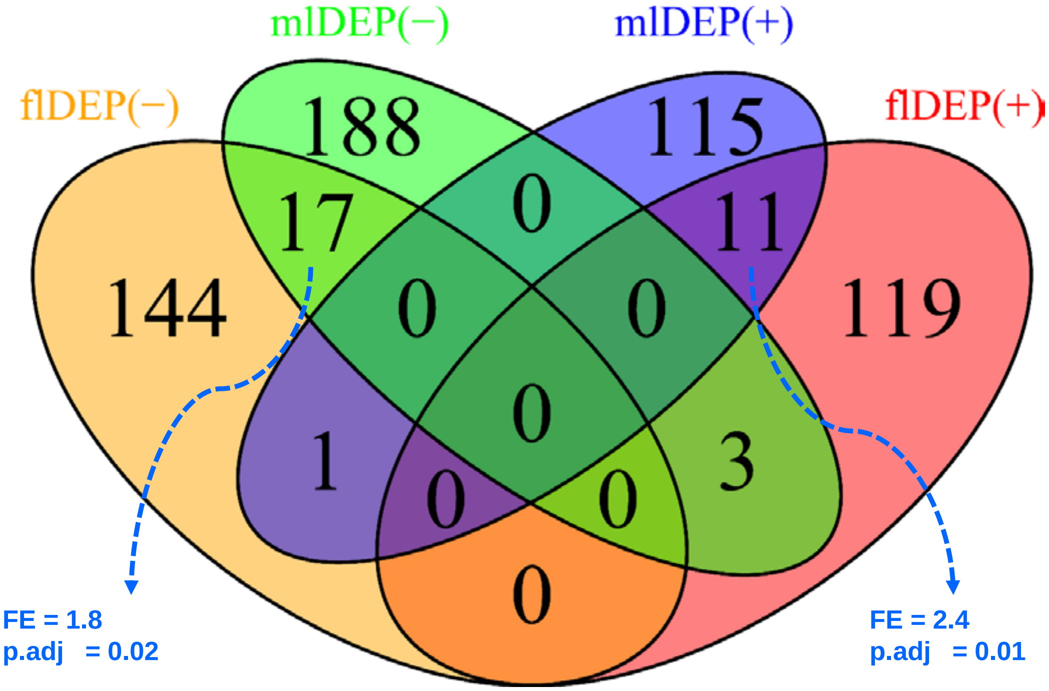

**B** **5xFAD-DUSP4vs5xFAD vs. 5xFAD-GFPvsWT in females**

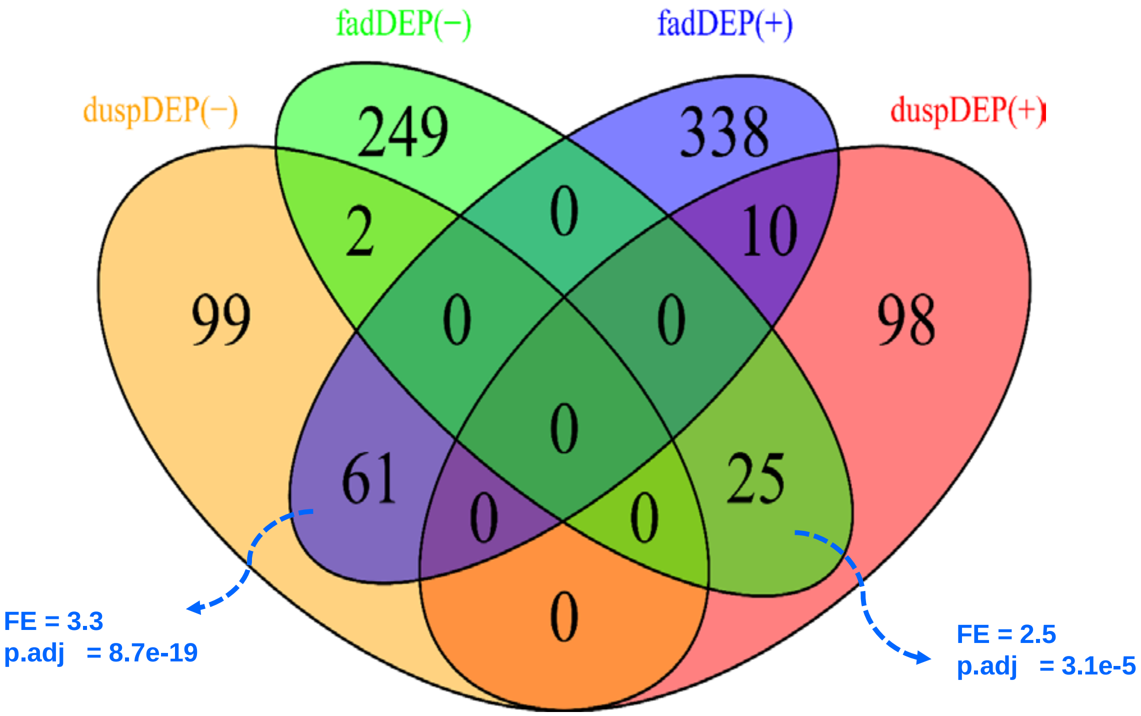

**Supplementary Figure 3. Venn diagrams highlighting the overlap of the DEP signatures.** (**A**) 5xFAD-DUSP4vs5xFAD in the male vs female mice. mlDEP(-) and mlDEP(+), are down- and up-regulated DEPs in males. flDEP(-) and flDEP(+), are down- and up-regulated DEPs in females. (**B**) Between 5xFAD-DUSP4vs5xFAD and 5xFADvsWT in the female mice. fadDEP(-) and fadDEP(+) are down- and up-regulated DEPs in 5XFAD. duspDEP(-) and duspDEP(+) are down- and up-regulated DEPs in DUSP4 OE. FE, fold enrichment. p.adj, BH-adjusted p value.

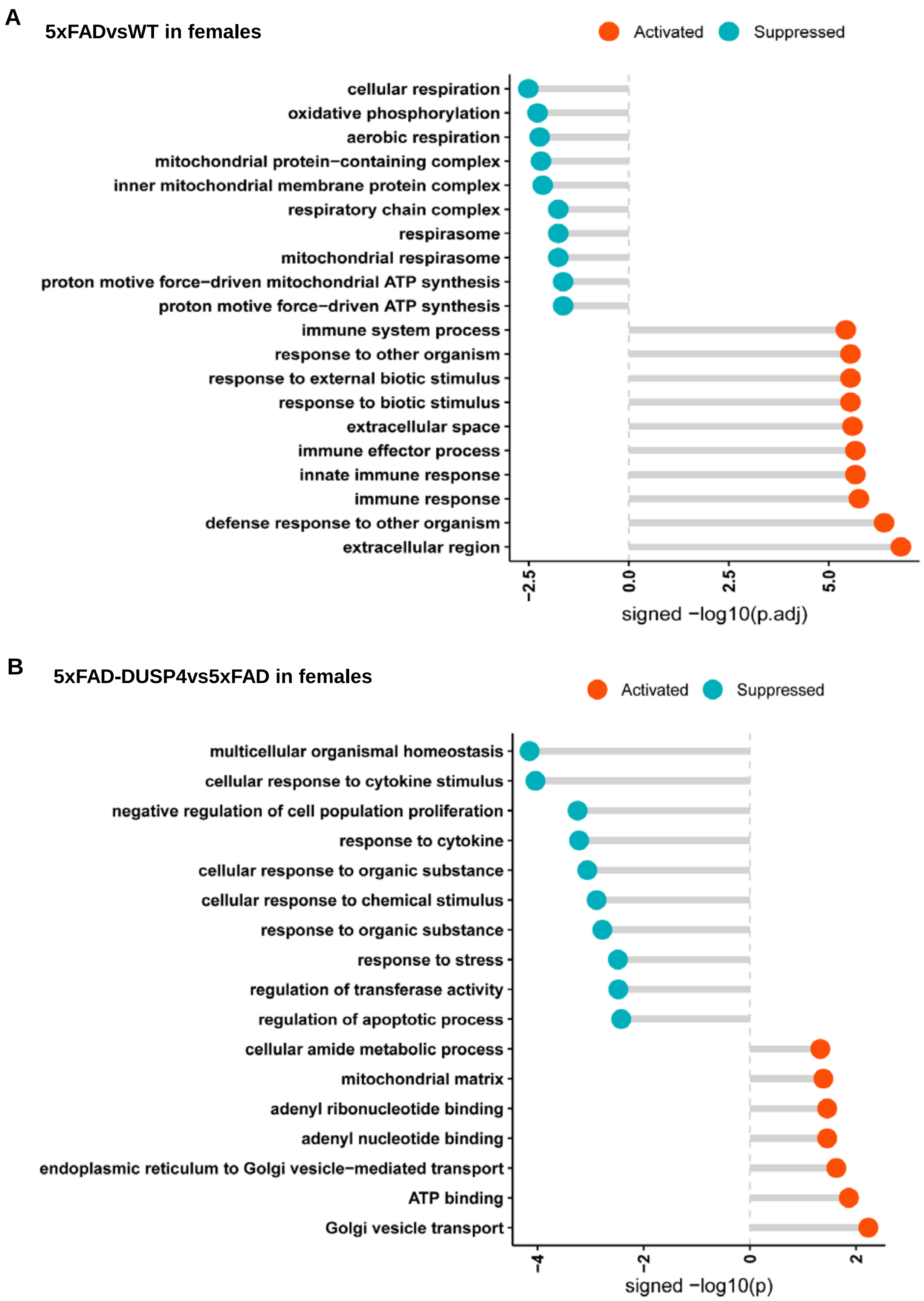
A
5xFADvsWT in females
Activated
Suppressed
cellular respiration
oxidative phosphorylation
aerobic respiration
mitochondrial protein−containing complex
inner mitochondrial membrane protein complex
respiratory chain complex
respirasome
mitochondrial respirasome
proton motive force−driven mitochondrial ATP synthesis
proton motive force−driven ATP synthesis
immune system process
response to other organism
response to external biotic stimulus
response to biotic stimulus
extracellular space
immune effector process
innate immune response
immune response
defense response to other organism
extracellular region
−2.5
0.0
2.5
5.0
signed −log10(p.adj)
B
5xFAD-DUSP4vs5xFAD in females
Activated
Suppressed
multicellular organismal homeostasis
cellular response to cytokine stimulus
negative regulation of cell population proliferation
response to cytokine
cellular response to organic substance
cellular response to chemical stimulus
response to organic substance
response to stress
regulation of transferase activity
regulation of apoptotic process
cellular amide metabolic process
mitochondrial matrix
adenyl ribonucleotide binding
adenyl nucleotide binding
endoplasmic reticulum to Golgi vesicle−mediated transport
ATP binding
Golgi vesicle transport
−4
−2
0
2
signed −log10(p)

**Supplementary Figure 4. GO enrichment analysis on the DEP signatures.** (**A**) 5xFADvsWT in female mice. (B) 5xFAD-DUSP4vs5xFAD in female mice. GO terms are grouped by activated (in red) or suppressed (in turquoise). x-axis, -log10(p.adj)(**A**) or -log10(p) (**B**) split by enrichment groups, activated (positive) vs. suppressed (negative). y-axis, GO terms.

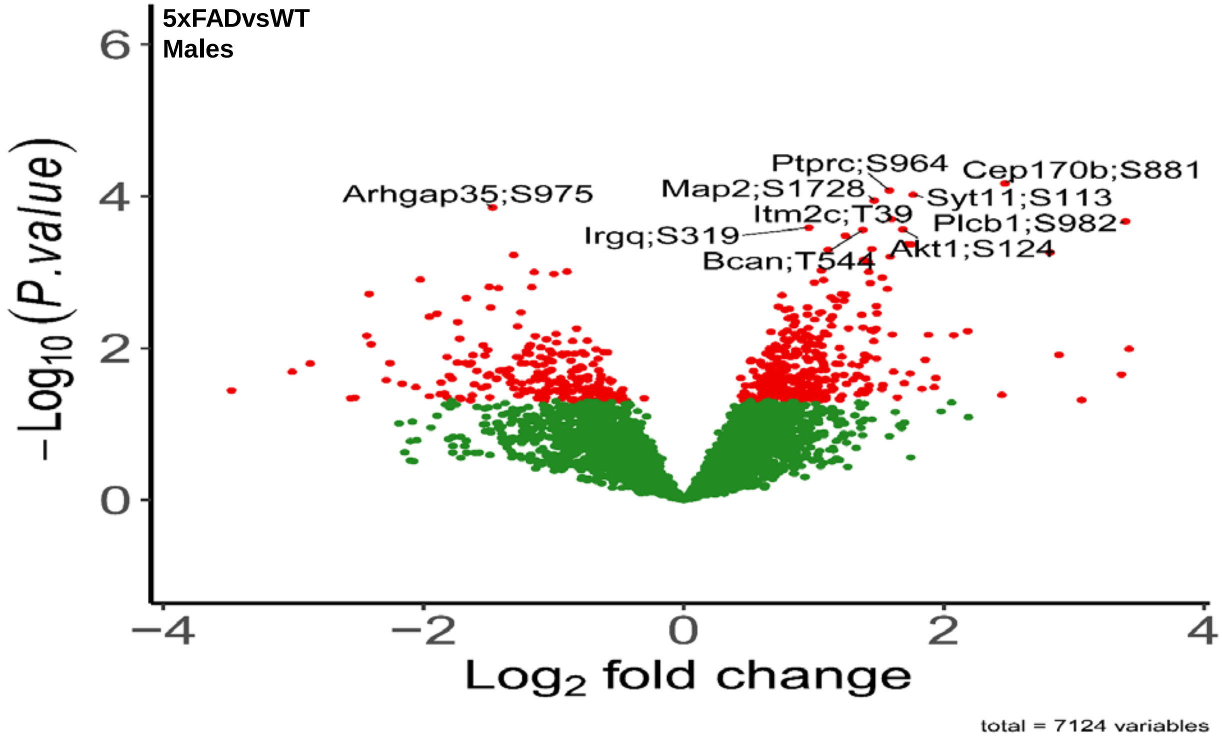
5xFADvsWT
Males
Ptprc;S964
Cep170b;S881
Map2;S1728
Syt11;S113
Arhgap35;S975
Itm2c;T39
Plcb1;S982
Irgq;S319
Akt1;S124
Bcan;T544
$-Log_{10}(P.value)$
$Log_2$ fold change
total = 7124 variables

B

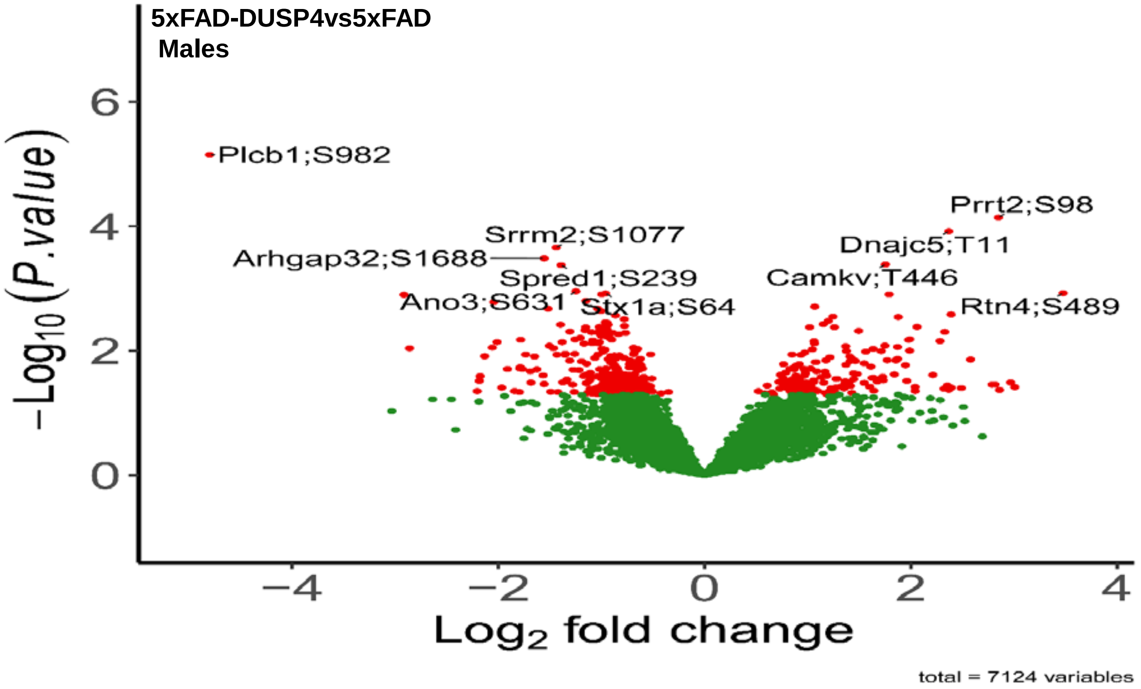
5xFAD-DUSP4vs5xFAD
Males
Plcb1;S982
Prrt2;S98
Srrm2;S1077
Dnajc5;T11
Arhgap32;S1688
Spred1;S239
Camkv;T446
Ano3;S631
Stx1a;S64
Rtn4;S489
$-Log_{10}(P.value)$
$Log_2$ fold change
total = 7124 variables

164

**Supplementary Figure 5. Volcano plots for DEPTM analysis.** (**A**) The DEPTMs in 5xFADvsWT in the male mice. (**B**) The DEPTMs in 5xFAD-DUSP4GFPvs5xFAD in the male mice. Each dot represents a PTM (phosphorylation) site. Highlighted are top-ranked PTMs (protein name and PTM site are connected by “;”). Dots in red are DEPTMs, whereas dots in green are not differentially expressed PTMs.

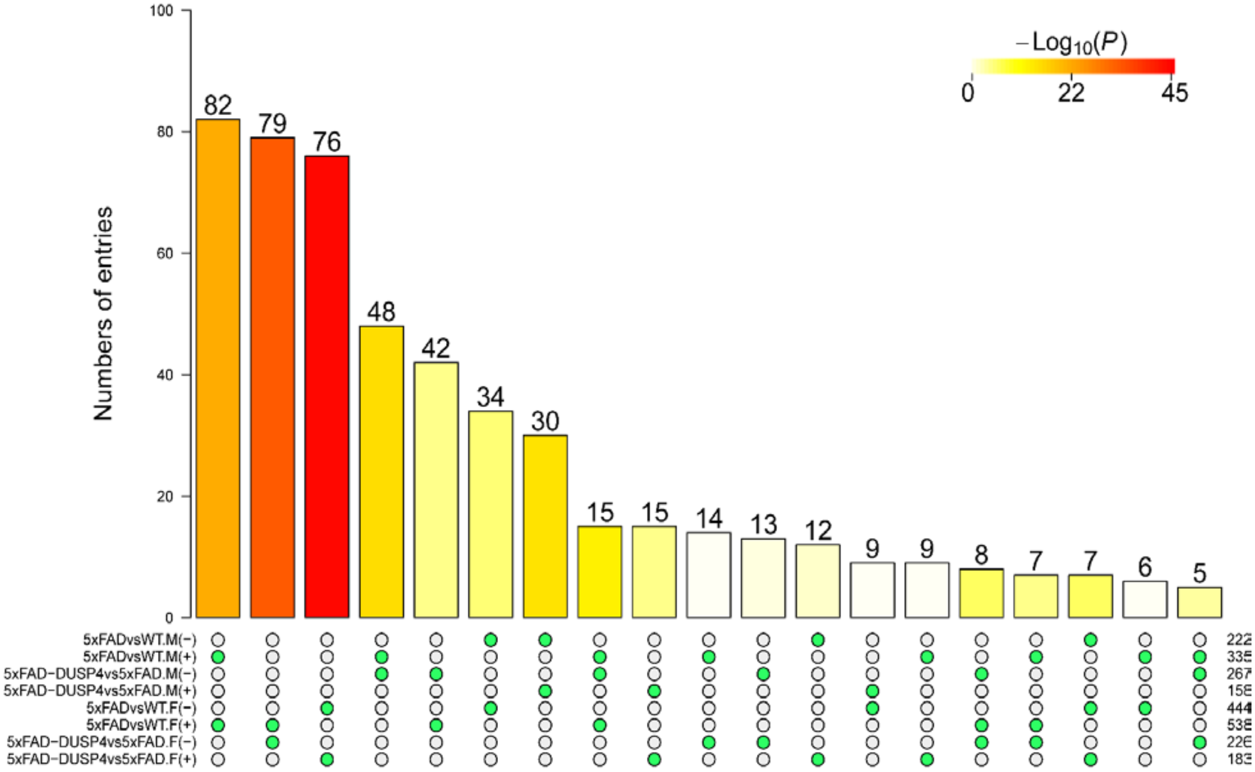

**Supplementary Figure 6. Summary of the overlap and significance of the DEPTM signatures via the superExactTest.** On the bottom of the bar graph, each row represents a DEPTM signature split by directionality in change in expression: - and +, down- and up-regulated, respectively. F and M, female and male, respectively. For example, 5xFADvsWT.M(-) stands for the down-regulated DEPTM signature in 5xFADvsWT in male mice. On the right are the numbers of the DEPTMs in each signature. Only shown here are the overlaps up to 3 groups. The number at the top of each bar is the number of DEPTMs overlapped among the groups (filled green circles)

across rows under each bar. The color in each bar is proportional to the significance of the overlap (-log10(P)).

**A**

**5xFAD-DUSP4vs5xFAD vs. 5xFAD-GFPvsWT in males**

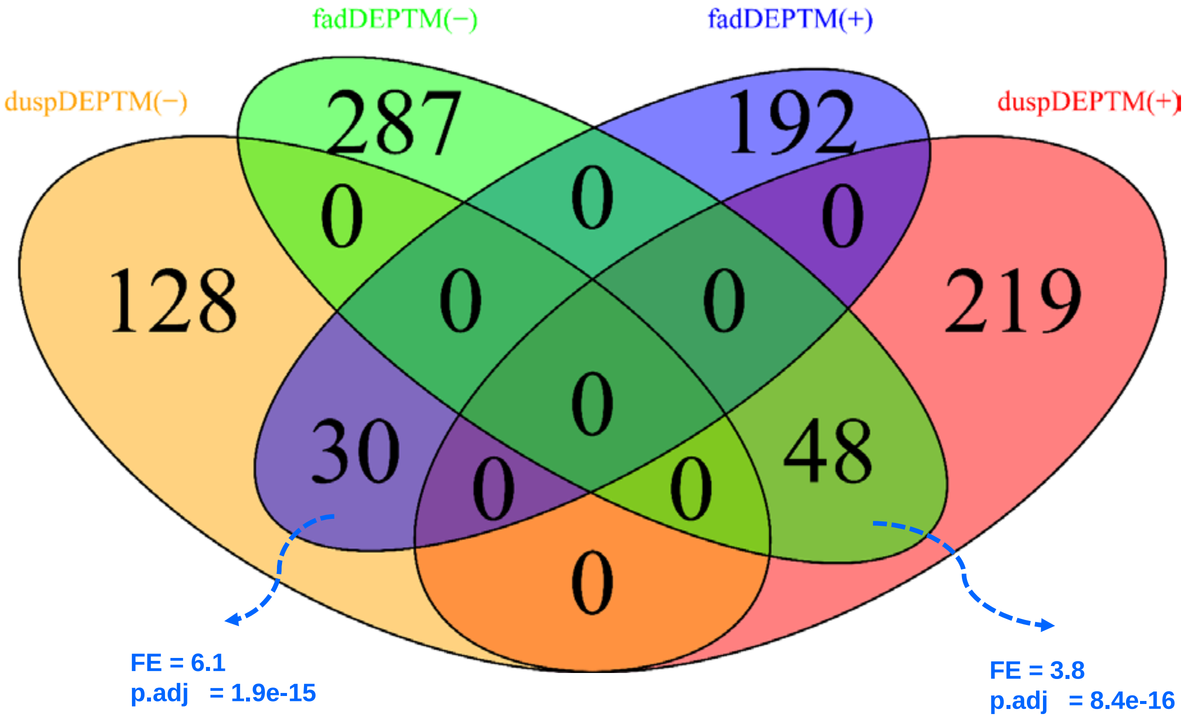

**B**

**5xFAD-DUSP4vs5xFAD in male vs female**

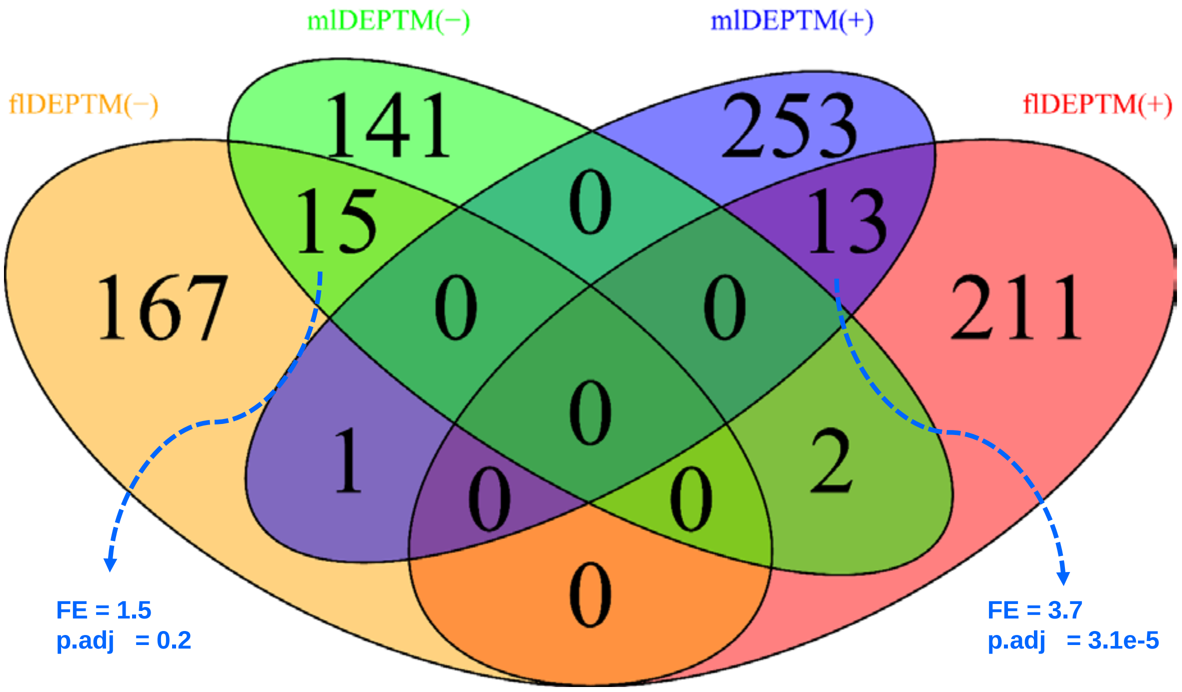

**Supplementary Figure 7. Venn diagrams highlighting for the overlap of DEPTM signatures.** (**A**) Between 5xFAD-DUSP4vs5xFAD and 5xFADvsWT in the male mice. fadDEPTM(-) and fadDEPTM(+) are down- and up-regulated DEPTMs in 5XFAD. duspDEPTM(-) and duspDEPTM(+) are down- and up-regulated DEPTMs in DUSP4 OE. (**B**) 5xFAD-DUSP4vs5xFAD in the male vs female mice. mlDEPTM(-) and mlDEPTM(+), are down- and up-regulated DEPTMs in male. flDEPTM(-) and flDEPTM(+), are down- and up-regulated DEPTMs in female. FE, fold enrichment. p.adj, BH-adjusted p value.

A

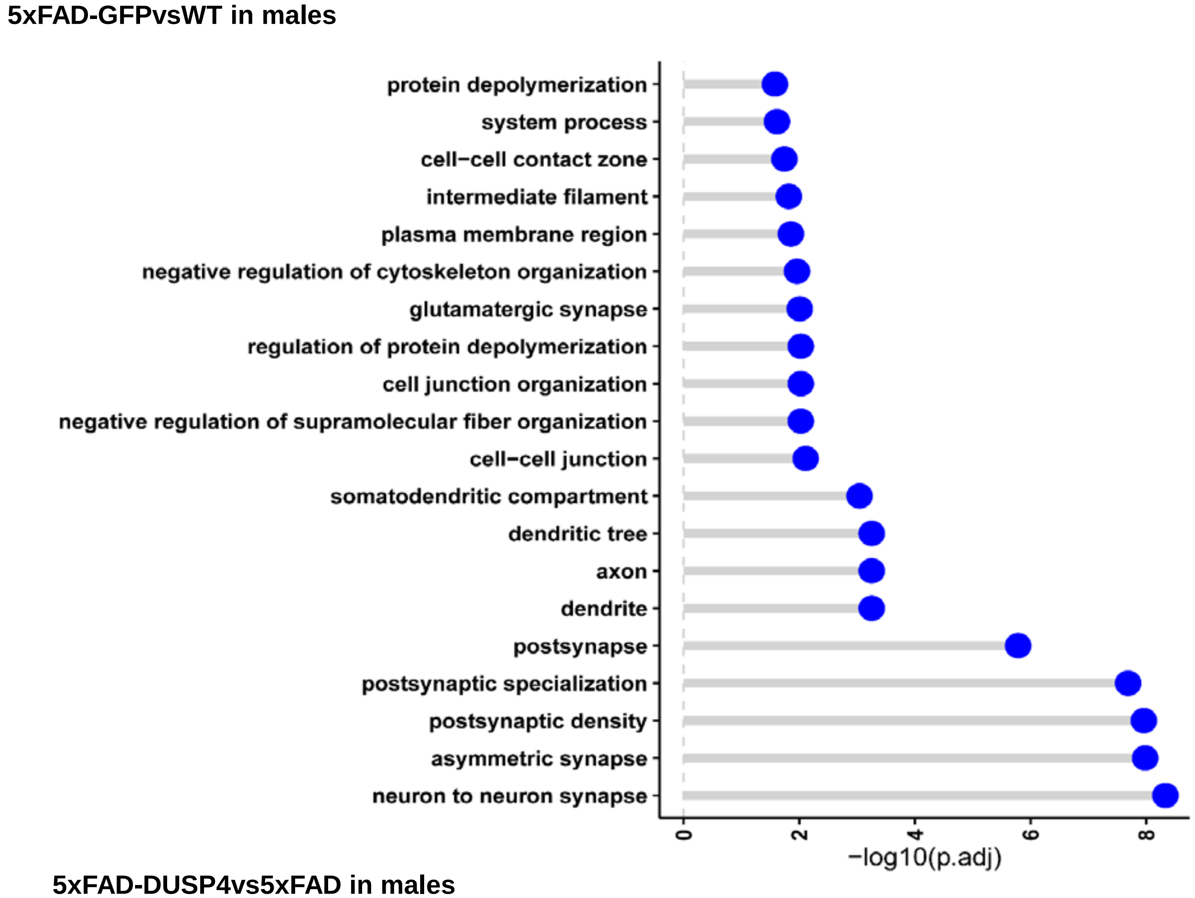

5xFAD-GFPvsWT in males
protein depolymerization
system process
cell−cell contact zone
intermediate filament
plasma membrane region
negative regulation of cytoskeleton organization
glutamatergic synapse
regulation of protein depolymerization
cell junction organization
negative regulation of supramolecular fiber organization
cell−cell junction
somatodendritic compartment
dendritic tree
axon
dendrite
postsynapse
postsynaptic specialization
postsynaptic density
asymmetric synapse
neuron to neuron synapse
0
2
4
6
8
−log10(p.adj)

B

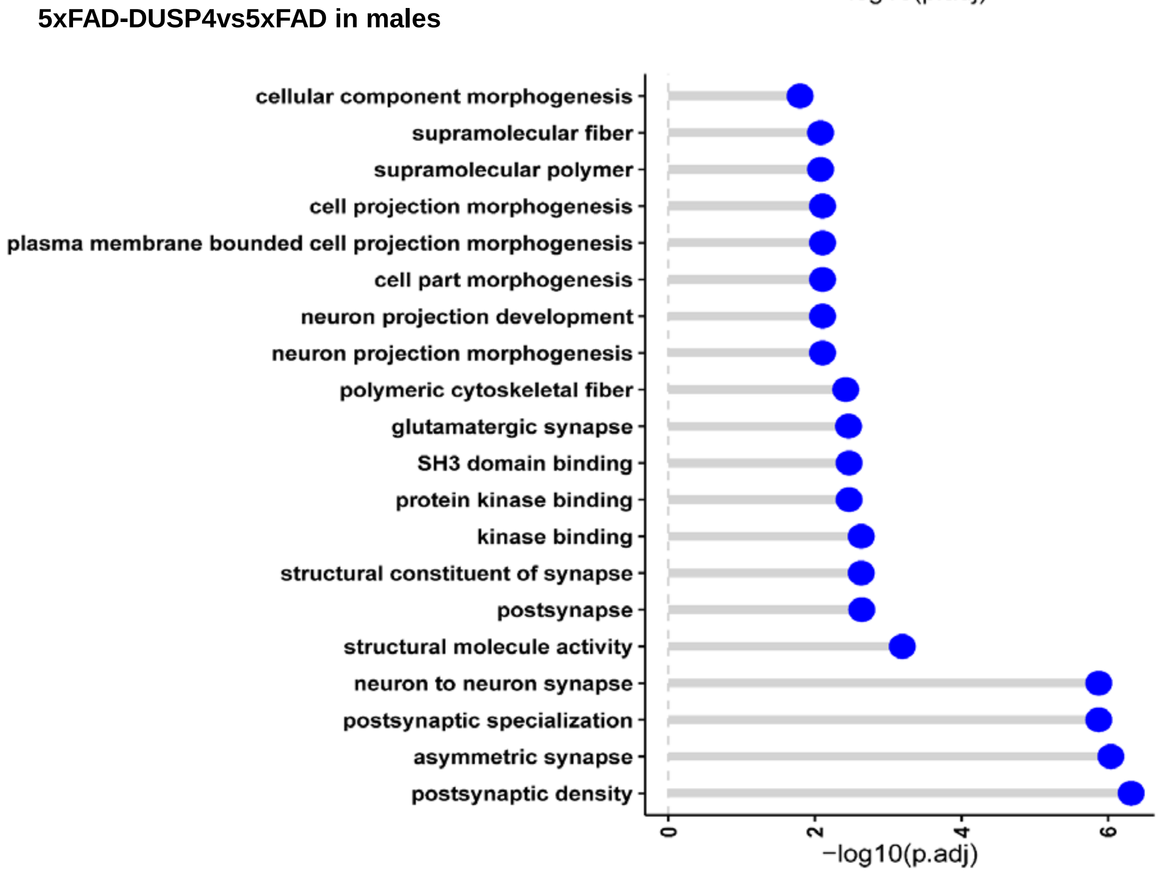

5xFAD-DUSP4vs5xFAD in males
cellular component morphogenesis
supramolecular fiber
supramolecular polymer
cell projection morphogenesis
plasma membrane bounded cell projection morphogenesis
cell part morphogenesis
neuron projection development
neuron projection morphogenesis
polymeric cytoskeletal fiber
glutamatergic synapse
SH3 domain binding
protein kinase binding
kinase binding
structural constituent of synapse
postsynapse
structural molecule activity
neuron to neuron synapse
postsynaptic specialization
asymmetric synapse
postsynaptic density
0
2
4
6
−log10(p.adj)

**Supplementary Figure 8. GO enrichment analysis on the DEPTM signatures.** (**A**) 5xFADvsWT in male mice. (B) 5xFAD-DUSP4vs5xFAD in male mice. DEPTMs were collapsed across each protein. y-axis, GO terms. x-axis, -log10(p.adj).

**A**

**5xFAD-DUSP4vs5xFAD in males**

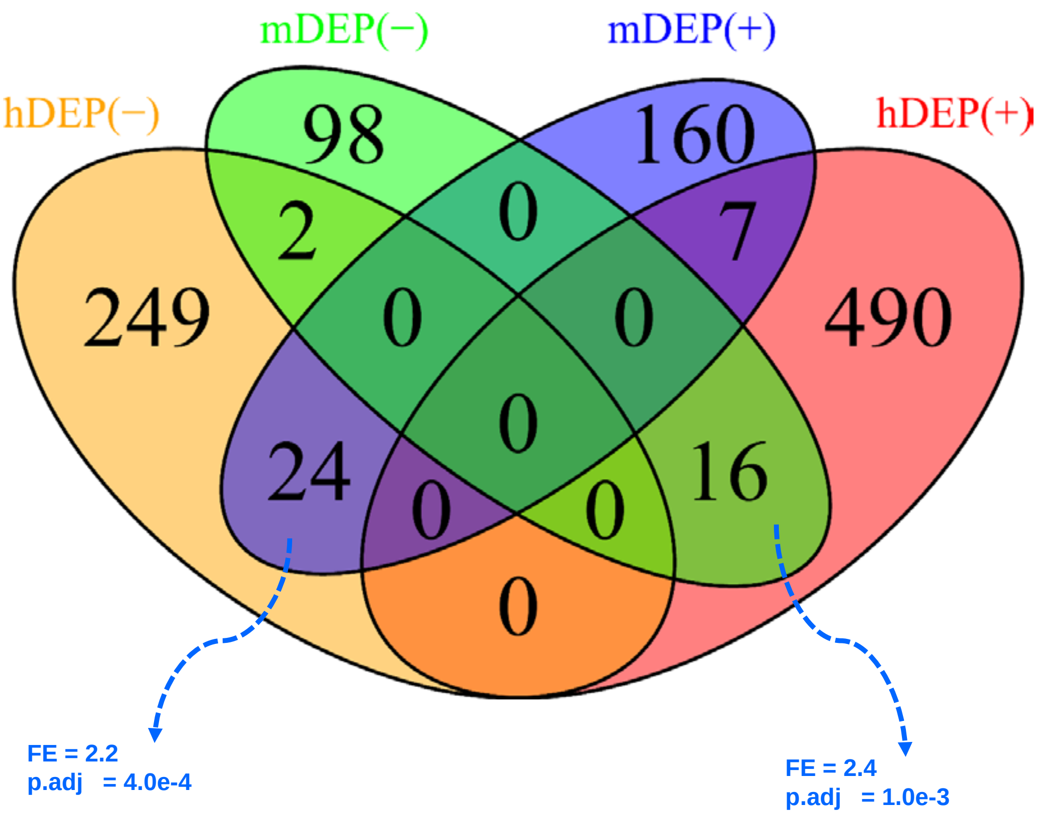

**B**

**5xFAD-DUSP4vs5xFAD in females**

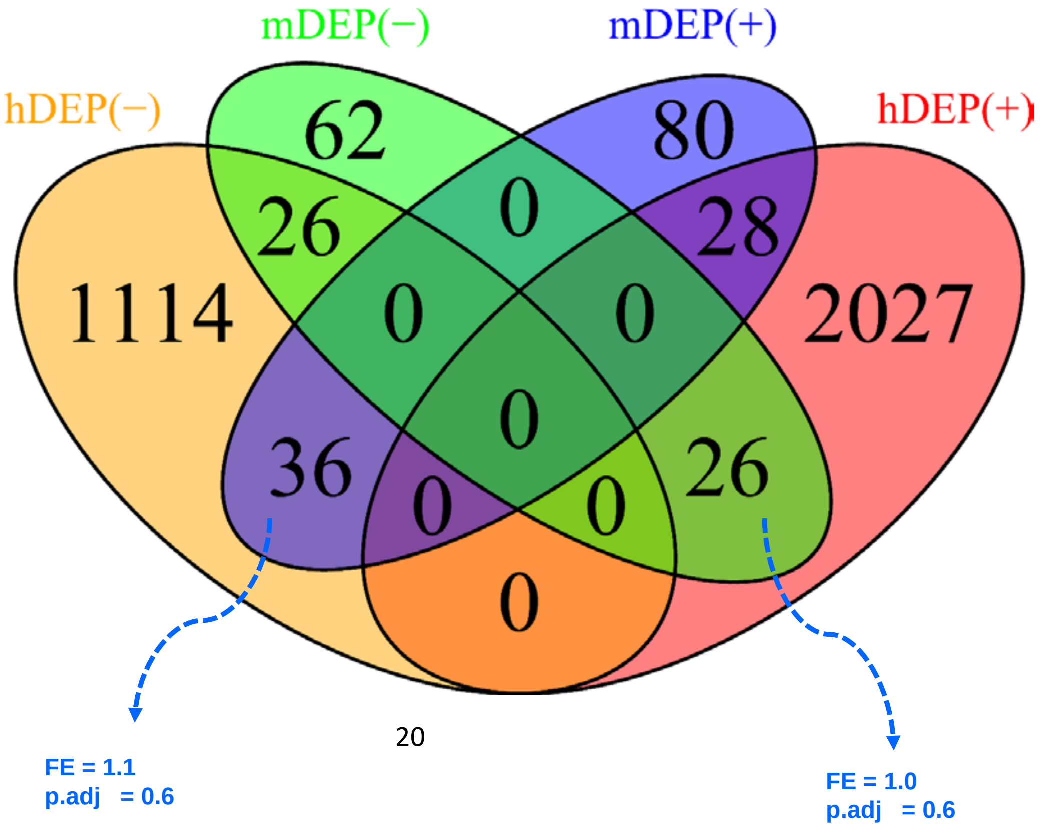

**Supplementary Figure 9. Venn diagrams highlighting for the overlap of the DEP signatures between human and mouse.** (**A**) Venn diagram revealing the overlap between the male mouse DEPs in 5xFAD-DUSP4vs5xFAD and the human male-specific DEPs in AD vs NL. (**B**) Venn diagram showing the overlap between the female mouse DEPs in 5xFAD-DUSP4vs5xFAD and the human female-specific DEPs in AD vs NL. In (A) and (B), mDEP(+) and mDEP(-), stand for up- and down-regulated DEPs in mice, respectively, whereas hDEP(+) and hDEP(-), stand for up- and down-regulated DEPs in human, respectively.

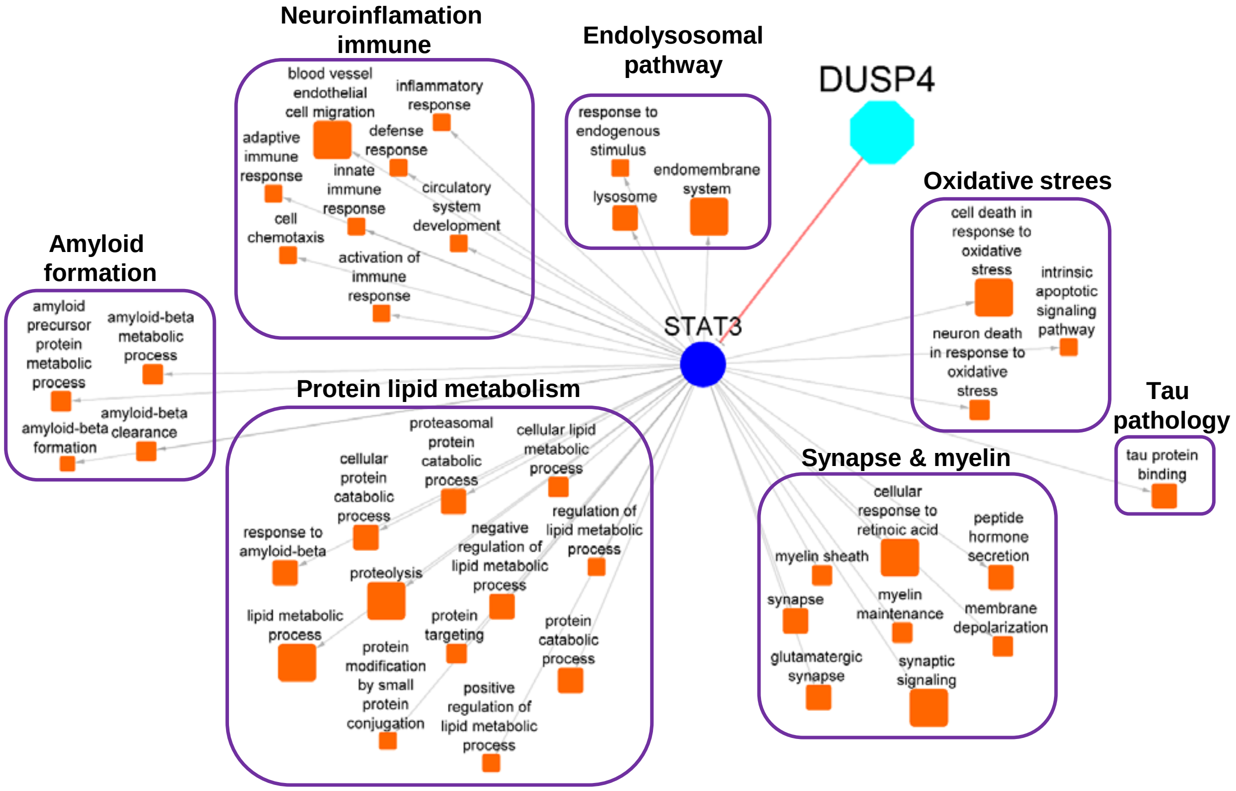

**Supplementary Figure 10. The DUSP4-STAT3 protein signal map.** The protein and protein interaction (PPI) subnetwork of STAT3 was obtained by querying the global PPI network[7] (kindly gift from Dr. Feixiong Cheng) up to 1 undirected walk. The STAT3 protein subnetwork was overlapped with the AD-related GO pathways[8,9] for enrichment, and thereby the DUSP4-STAT3 signal map was constructed (see Methods). Each filled box stands for a GO term, whose size is proportional to its enrichment for the STAT3 subnetwork with a GO term. Large unfilled box are for the parent categories of GO terms in AD.

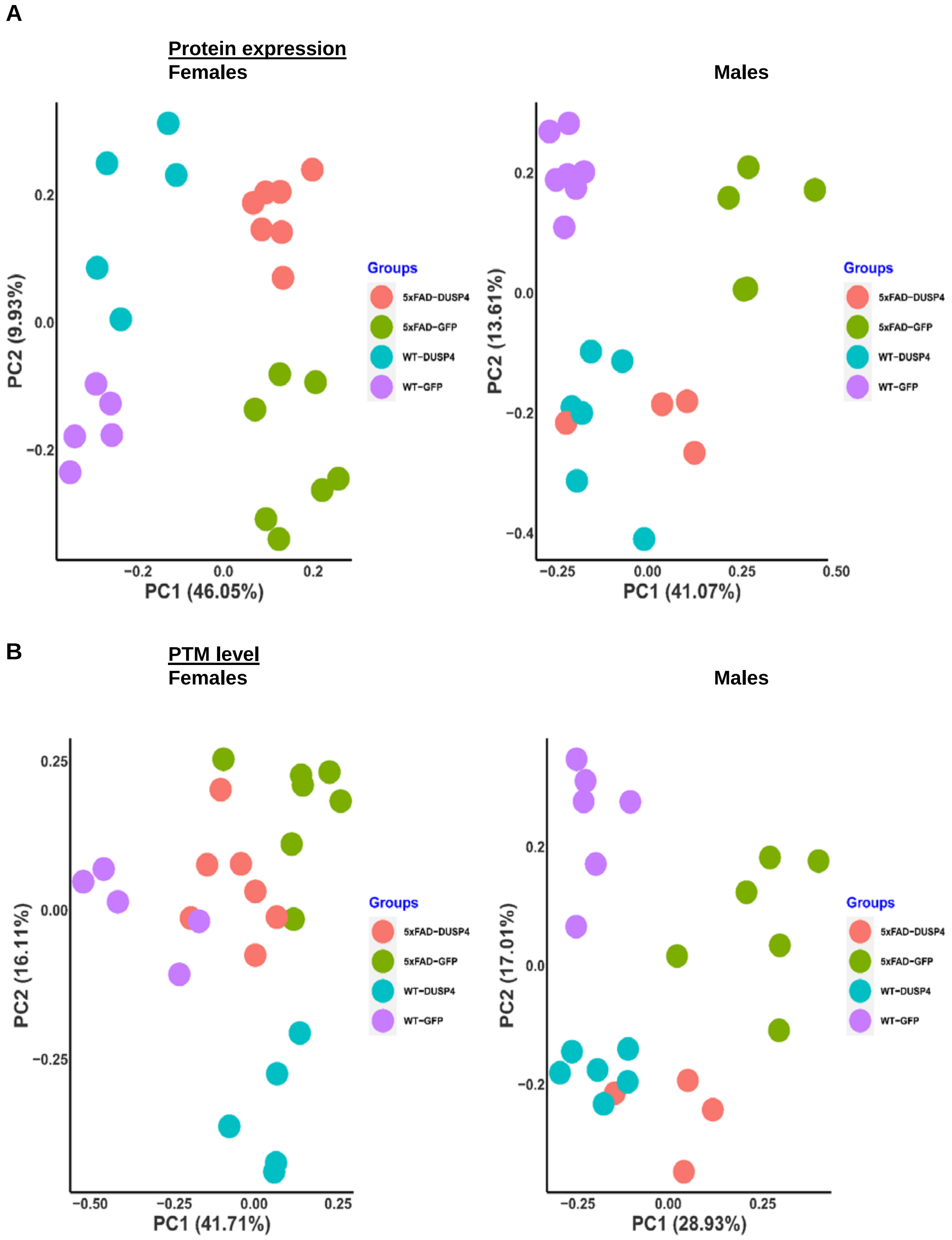
A
Protein expression
Females
Males
PC1 (46.05%)
PC2 (9.93%)
PC1 (41.07%)
PC2 (13.61%)
Groups
5xFAD-DUSP4
5xFAD-GFP
WT-DUSP4
WT-GFP
B
PTM level
Females
Males
PC1 (41.71%)
PC2 (16.11%)
PC1 (28.93%)
PC2 (17.01%)

**Supplementary Figure 11. PCA analysis of the protein and phosphoprotein expression.** (**A**) The protein expression in the male (right panel) and female mice (left panel). (**B**) The phosphoprotein expression in the male (right panel) and female mice (left panel). Overall, the 4 experimental groups of mice can be classified by their genotype groups via PCA.

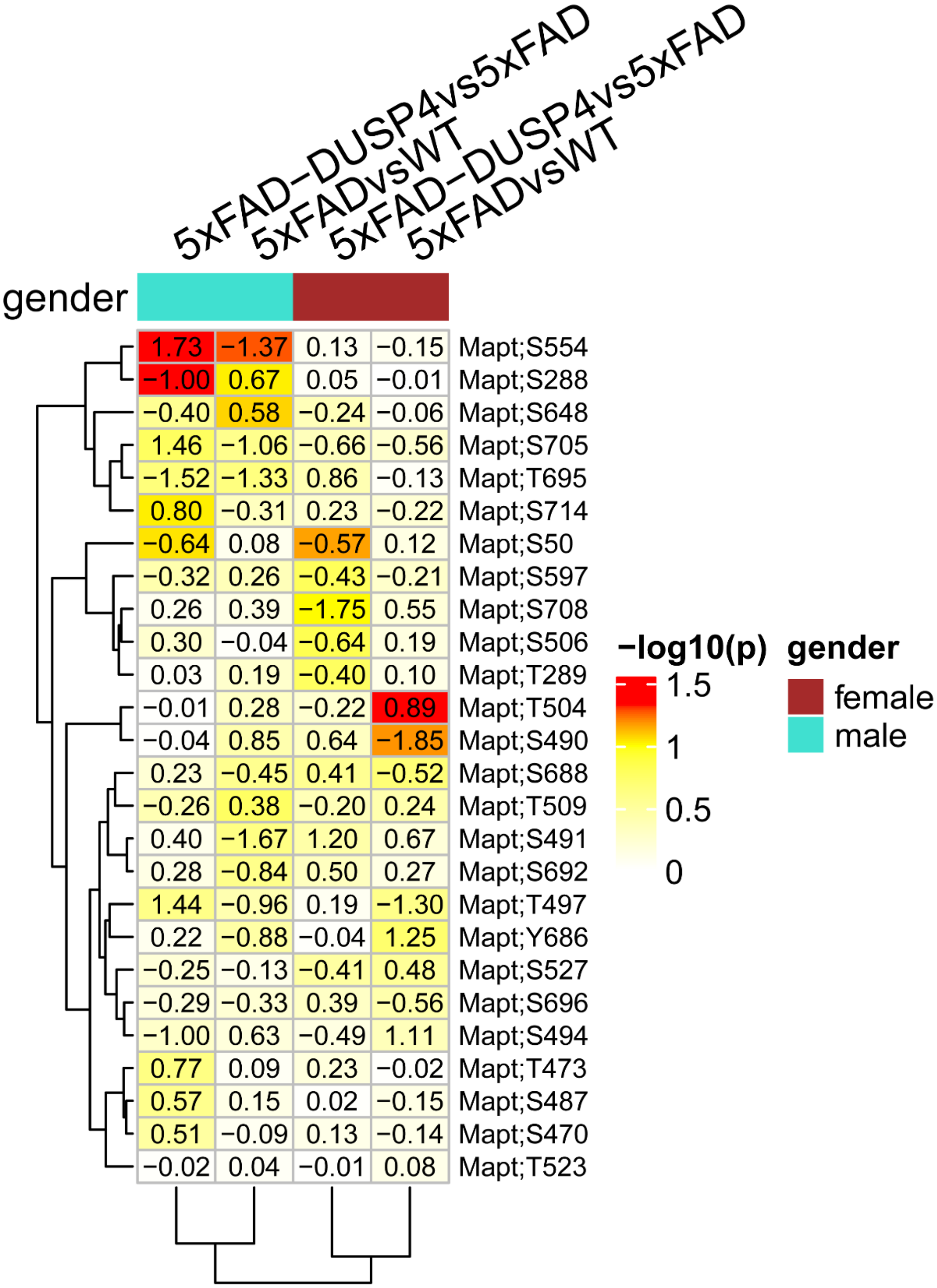
5xFAD−DUSP4vs5xFAD
5xFADvsWT
5xFAD−DUSP4vs5xFAD
5xFADvsWT
gender
1.73 −1.37 0.13 −0.15 Mapt;S554
−1.00 0.67 0.05 −0.01 Mapt;S288
−0.40 0.58 −0.24 −0.06 Mapt;S648
1.46 −1.06 −0.66 −0.56 Mapt;S705
−1.52 −1.33 0.86 −0.13 Mapt;T695
0.80 −0.31 0.23 −0.22 Mapt;S714
−0.64 0.08 −0.57 0.12 Mapt;S50
−0.32 0.26 −0.43 −0.21 Mapt;S597
0.26 0.39 −1.75 0.55 Mapt;S708
0.30 −0.04 −0.64 0.19 Mapt;S506
0.03 0.19 −0.40 0.10 Mapt;T289
−0.01 0.28 −0.22 0.89 Mapt;T504
−0.04 0.85 0.64 −1.85 Mapt;S490
0.23 −0.45 0.41 −0.52 Mapt;S688
−0.26 0.38 −0.20 0.24 Mapt;T509
0.40 −1.67 1.20 0.67 Mapt;S491
0.28 −0.84 0.50 0.27 Mapt;S692
1.44 −0.96 0.19 −1.30 Mapt;T497
0.22 −0.88 −0.04 1.25 Mapt;Y686
−0.25 −0.13 −0.41 0.48 Mapt;S527
−0.29 −0.33 0.39 −0.56 Mapt;S696
−1.00 0.63 −0.49 1.11 Mapt;S494
0.77 0.09 0.23 −0.02 Mapt;T473
0.57 0.15 0.02 −0.15 Mapt;S487
0.51 −0.09 0.13 −0.14 Mapt;S470
−0.02 0.04 −0.01 0.08 Mapt;T523
−log10(p)
1.5
1
0.5
0
gender
female
male

**Supplementary Figure 12. Heatmap showing the significance and change in expression of PTMs in the tau protein (Mapt gene).** The color intensity denotes and is proportional to the –log10(p), where p is the statistic p value in each comparison. Highlighted numbers are the fold change in expression in each comparison.

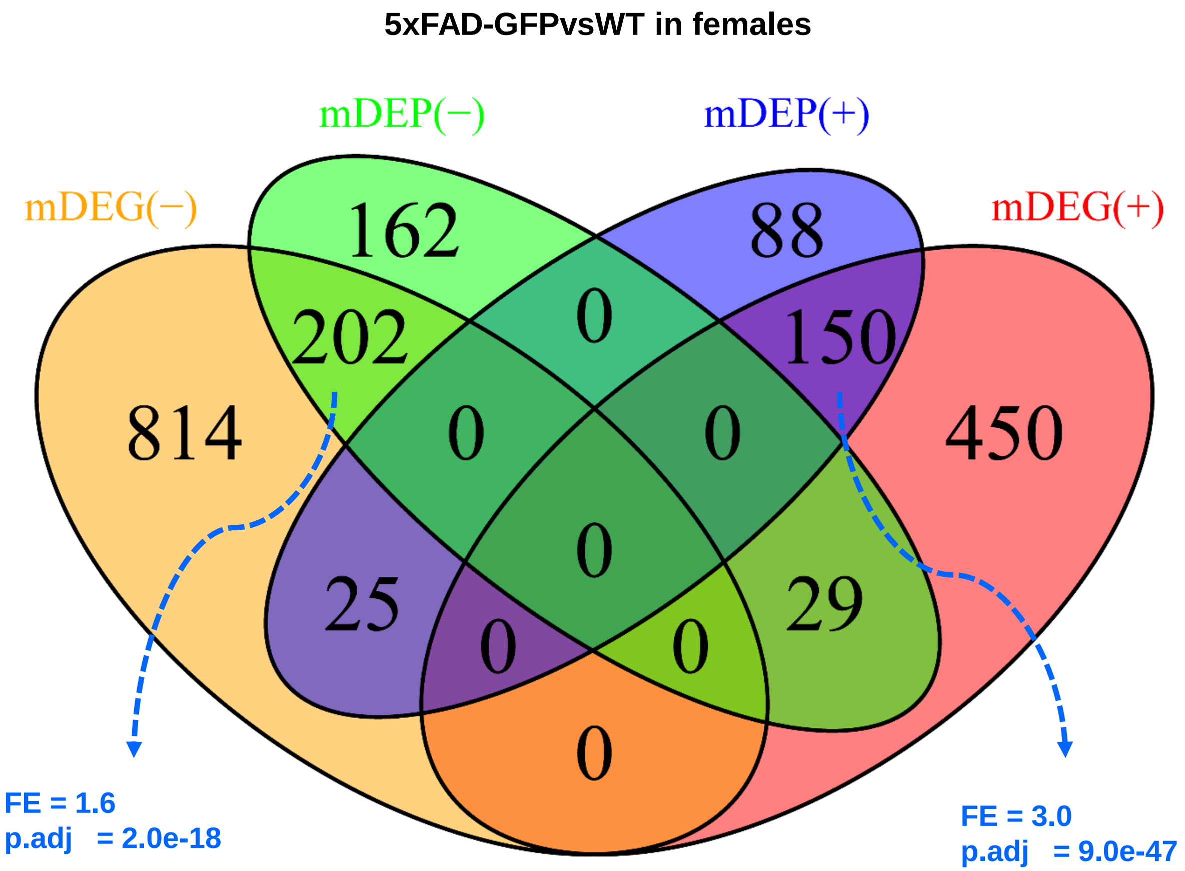

**Supplementary Figure 13. Venn diagram highlighting for the overlap between DEP and DEG signatures for 5xFADvsWT.** mDEP(-) and mDEP(+), are down- and up-regulated DEPs in females, respectively. mDEG(-) and mDEG(+), are down- and up-regulated DEGs in females, respectively. FE, fold enrichment. p.adj, BH-adjusted p value.

**A**

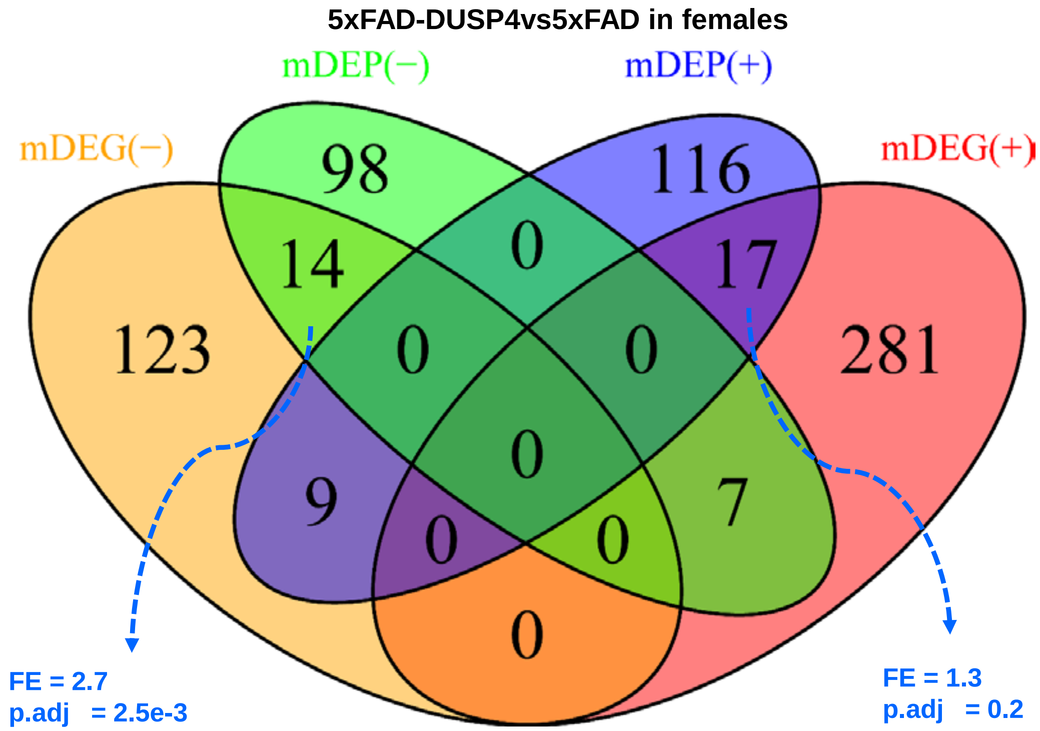
5xFAD-DUSP4vs5xFAD in females
mDEP(−)
mDEP(+)
mDEG(−)
mDEG(+)
98
116
14
0
17
123
0
0
281
0
9
7
0
0
0
FE = 2.7
p.adj = 2.5e-3
FE = 1.3
p.adj = 0.2

**B**

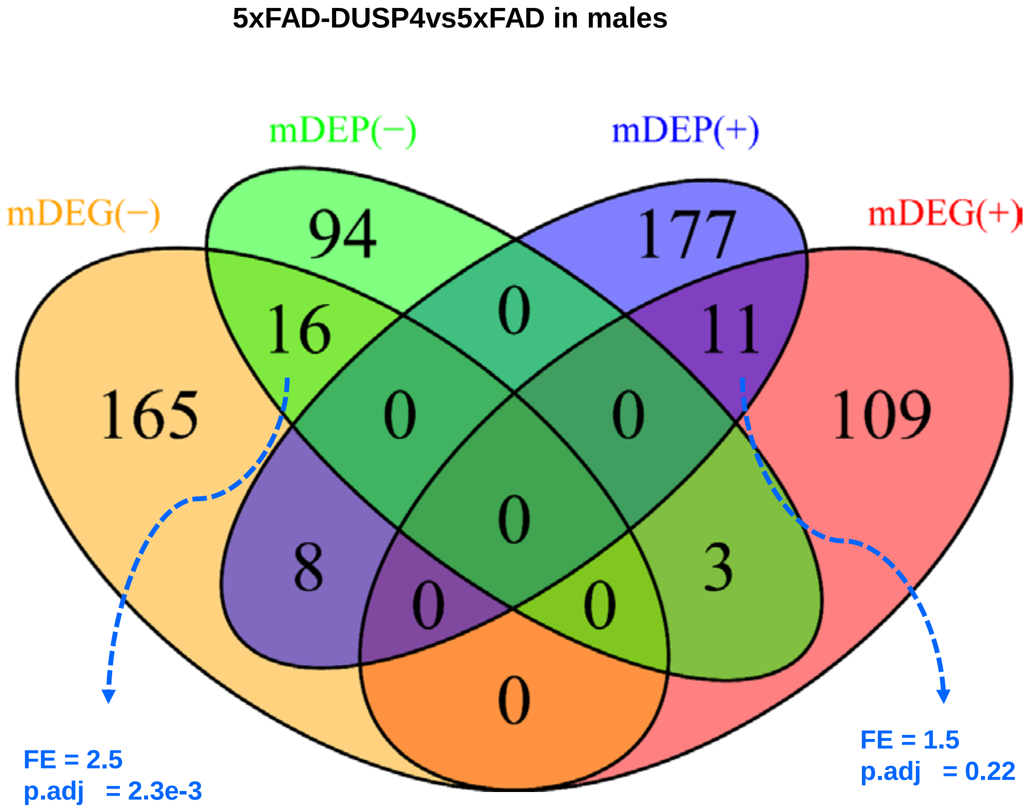
5xFAD-DUSP4vs5xFAD in males
mDEP(−)
mDEP(+)
mDEG(−)
mDEG(+)
94
177
16
0
11
165
0
0
109
0
8
3
0
0
0
FE = 2.5
p.adj = 2.3e-3
FE = 1.5
p.adj = 0.22

**Supplementary Figure 14. Venn diagrams highlighting for the overlap between DEP and DEG signatures for 5xFAD-DUSP4vs5xFAD.** (**A**) female mice; (B) male mice. mDEP(-) and mDEP(+), are down- and up-regulated DEPs, respectively. mDEG(-) and mDEG(+), are down- and up-regulated DEGs, respectively. FE, fold enrichment. p.adj, BH-adjusted p value.

### Supplementary Data

Supplementary.data1.xlsx

Supplementary.data2.xlsx

Supplementary.data3.xlsx

Supplementary.data4.xlsx
